# Effects of homoeologous exchange on gene expression and alternative splicing in a newly formed allotetraploid wheat

**DOI:** 10.1101/2021.12.28.474319

**Authors:** Zhibin Zhang, Hongwei Xun, Ruili Lv, Xiaowan Gou, Xintong Ma, Juzuo Li, Jing Zhao, Ning Li, Lei Gong, Bao Liu

## Abstract

Homoeologous exchange (HE) is a major mechanism generating post-polyploidization genetic variation with important evolutionary consequences. However, the direct impacts of HE without entangling with additional evolutionary forces on gene expression remains to be fully understood. Here, we analyzed high-throughput RNA-seq data of young leaves from individuals of a synthetic allotetraploid wheat (AADD), which contain variable numbers of HEs. We aimed to investigate if and to which extent HE directly impacts gene expression and alternative splicing (AS). We found that HE impacts expression of genes located within HE regions primarily via *cis*-acting dosage effect, which led to significant changes in the total expression of homoeolog pairs, especially for homoeologs whose original expression was biased. In parallel, HE influences expression of a large amount of genes residing in non-HE regions by *trans*-regulation leading to convergent expression of homoeolog pairs. Intriguingly, when taking into account of the original relative homoeolog expression states, homoeolog pairs under *trans*-effect are more prone to showing convergent response to HE whereas those under *cis*-effect trended to show subgenome-specific expression. Moreover, HE induced quantitative, largely individual-specific, changes of alternative splicing (AS) events. Like homoeologs expression, homoeo-AS events which related to *trans* effect were more responsive to HE. HE therefore exerts multifaceted immediate effects on gene expression and, to a less extent, also transcript diversity in nascent allopolyploidy.

## Introduction

Polyploidization or whole-genome duplication (WGD) is a major driving force in genome evolution and speciation throughout the evolutionary history of higher plants (**Wendel, 2000, Van de Peer *et al*., 2009, Jiao *et al*., 2011)**. After merging and doubling two or more divergent genomes in the same nucleus, allopolyploidy could cause rapid genetic and epigenetic re-programing, i.e., “genomic shock” **(McClintock, 1984, Adams, 2007, Madlung and Wendel, 2013, Diez *et al*., 2014, Ding and Chen, 2018)**, which may provide substrates for natural selection and hence enable successful evolution of allopolyploid populations and facilitates speciation **(Wendel, 2000, Comai, 2005, Crow and Wagner, 2005, Otto, 2007, Feldman *et al*., 2012, Schoenfelder and Fox, 2015)**.

As a mechanism unique to allopolyploidy, homoeologs exchange (HE) refers to the exchange of chromosome segments between homoeologous chromosomes with high sequence similarity during meiosis **(Hollister, 2015, Hurgobin *et al*., 2018, Lloyd *et al*., 2018)**. In synthetic and young allopolyploids, HE provides a mechanism for rapid genomic variation **(Gaeta and Chris Pires, 2010, Hollister, 2015, Mercier *et al*., 2015, Bomblies *et al*., 2016)**, which may help circumvent the severe genetic bottleneck intrinsic to allopolyploidy speciation (**Wu *et al*., 2021**). HE was found to be widely present in many young allopolyploid crops, such as *Brassica* **(Osborn *et al*., 2003, Pires *et al*., 2004, Xiong *et al*., 2011, Chalhoub *et al*., 2014, Stein *et al*., 2017, Lloyd *et al*., 2018)**, *Musa* **(Wang *et al*., 2019)**, *Fragaria* **(Edger et al., 2019)**, *Coffea* **(Lashermes et al., 2014)**, *Arachisc* **(Bertioli et al., 2019)** and occasionally in wheat **(He *et al*., 2017)**. This suggests HE-underpinned genetic variants have been selected for during domestication. Indeed, it was documented that HE confers high adaptability and produces a wide range of physiological and phenotypic variations desired in crops **(Pires *et al*., 2004, Stein *et al*., 2017, Raman *et al*., 2021, Shi *et al*., 2021)**. It was also suggested that under natural settings HE plays important parts in generating evolutional novelty, accelerating diploidization process and facilitating speciation **(Edger *et al*., 2018, Mason and Wendel, 2020)**.

At the molecular level, HE-induced chromosome segment reshuffling may cause or contribute to dramatic genomic, epigenomic and transcriptomic changes. In genome aspect, pan-genome analysis of *B. napus* indicates that ~38% of genes with presence/absence variation (PAV) pattern is associated with HE, and such gene set is involved in essential agronomic traits including flowering time, disease resistance, acyl lipid metabolism and glucosinolate metabolism **(Hurgobin *et al*., 2018)**. In epigenome aspect, HE was shown to sustain or even further enhance WGD-induced DNA methylation repatterning that is correlated with gene expression **(Li *et al*., 2019)**. In gene expression aspect, a recent study showed that the magnitude of expression changes was positively correlated to the degree of allele copy number induced by HE (*cis* regions), that is, there is a dosage effect on gene expression; moreover, when homoeolog expression bias present, duplication of one homoeolog allele cannot compensate for the loss of the other homoeolog allele, leading to inconsistency (and hence diversity) of the overall expression of homoeolog pairs after HE occurrence **(Lloyd *et al*., 2018)**. In addition, a study on synthetic allopolyploid rice unraveled that the combined effects of WGD and HE contributed to large-scale changes of gene alternative splicing (AS) patterns **(Zhang *et al*., 2019)**. However, these studies have primarily focused on HE-induced *cis* effects rather than *trans* regulation **(Lloyd’s work)** or did not distinguish the confounding effects of WGD and HE on RNA-transcription **(Zhang’s work)**. Therefore, it is necessary to systematically analyze how HE itself impacts on the atlas of RNA-transcription (including gene expression and AS) through the combination of *cis* and *trans* effects.

Common wheat (*Triticum aestivum*) with three distinct subgenomes, A (from *T. urartu*), B (from an unknown species related to *Aegilops speltoides*, genome SS), and D (from *Ae. tauschii*), was formed via two rounds of allopolyploidization events and provides a suitable system to study WGD-related genome evolution **(Matsuoka, 2011, Feldman and Levy, 2012, Marcussen *et al*., 2014)**. However, due to action of homoeologous pairing control genes, such as *Ph1and Ph2*, HE is rare in naturally formed tetraploid wheat and hexaploid common wheat **(Martínez-Pérez *et al*., 1999, Roberts *et al*., 1999, Martinez-Perez *et al*., 2001, Griffiths *et al*., 2006, Serra *et al*., 2021)**. In contrast, HEs are rampant in synthetic allotetraploids parented by the putative diploid progenitor species of common wheat, such as AADD and SSDD **(Zhang *et al*., 2013, Gou *et al*., 2018)**. Therefore, synthetic allopolyploid wheats enable assessing the direct effects of HE on genome and transcriptome changes and phenotypic diversity. Using synthetic tetraploid wheat AADD, we demonstrated that HE-induced intragenic recombination could generate novel, and potentially functional, transcripts, which provided a new aspect of the genetic consequences of HE **(Zhang *et al*., 2020)**. This also raises new questions: besides junction regions, does HE also impact the expression of genes located within HE and on non-HE genomic regions? Moreover, recent studies showed that polyploidization and domestication can generate qualitative (gain and/or loss) changes of AS events in natural wheat species **(Yu *et al*., 2020, Gao *et al*., 2021)**. So, is the direct effect of HE on AS in synthetic allopolyploids similar to that seen in established allopolyploid species that experienced natural selection and domestication?

In this study, we performed comprehensive analyses of gene expression and AS landscape in selfed progenies of a synthetic allotetraploid wheat (AADD) that have individuals with variable numbers of HEs. Our results have provided new insights into the direct impacts of HE on genome-wide transcript abundance and diversity. We show that HE exerts its action via both *cis*- and *trans*-effects depending on physical relationship of the genes with HE, as well as the influence of original expression states between members of a given homoeolog pair.

## Results

### Expression pattern of genes located in HE-related regions

Whether gene expression changes scale with gene copy number is a major reflection of the effects of HE on transcriptome variation in allopolyploids. We observed that the transcriptome atlas matched well with corresponding DNA copy numbers along all chromosomes, namely, genes located within HE regions (HE-related genes) showed proportional increase (duplication of DNA copies) or decrease (deletion of DNA copies) in expression levels (**Figure 1a and 1b**). Specifically, we assessed gene expression of euploid tetraploid individuals that contain various numbers of HEs (G1-J) together with an individual (A) as a control that contains a single HE that is shared by all the individuals. As shown in **Figure S1**, expression levels of genes located within the HE regions (HE-related genes) were lineally proportional to gene copy numbers, indicating HE-related genes mainly show dosage effects on expression.

**Figure 1.**
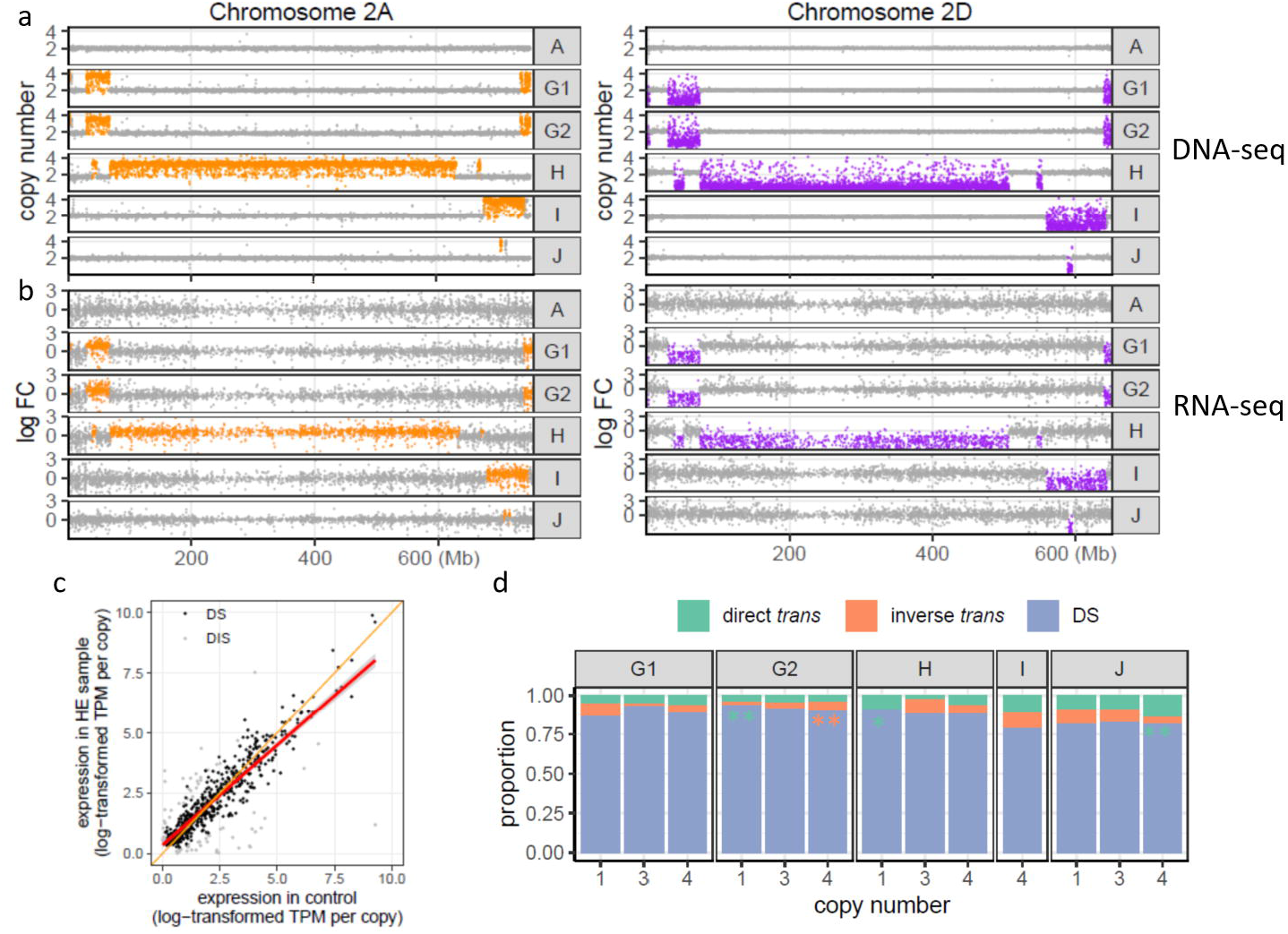
Gene expression pattern in HE regions. **(a)** DNA copy number and **(b)** relative expression plot along the chromosome 2A (left panel) and 2D (right panel), respectively. The copy numbers of tetraploid individuals in **(a)** were normalized based on *in silico* tetraploid, whereas the relative expression values in **(b)** were normalized based on the expression level (TPM) of *in silico* tetraploid. **(c)** An example of dosage-sensitive expression pattern of genes located on HE region in individual G1. The x-axis represents the expression level divided by the corresponding copy number (here is two) in the control plant, and the y-axis represents the expression level divided by the corresponding copy number (here is four) in G1. Black and grey dots indicate genes with dosage-sensitive (DS) and dosage-insensitive (DIS) expression, respectively. Red line represents the fitting curve based on real data, whereas the orange line indicates the theoretical dosage-sensitive trend line (y = x). **(d)** The proportion of DS, direct *trans* effect and inverse *trans* effect gene sets in HE regions of different copy numbers in each HE-containing individual. Green and orange asterisks represent proportion bias in genes with direct *trans* effect and with inverse *trans* effect, respectively.

To further quantify the contribution of dosage effects, size-factor-based differential expression analysis was performed to identify dosage-sensitive and -insensitive genes (**(Lloyd *et al*., 2018**); see Materials and Methods). In total, 78.7-92.0% HE-related genes (from 2,415 out of 3,068 genes in individual I to 4,049 out of 4,399 genes in individual G2) showed expression levels that are in strict concordance with corresponding DNA copy numbers (**Figure 1c and 1d; Figure S2**), i.e. dosage-sensitive (DS). The remaining genes are dosage insensitive, that is, their expression levels are constant irrespective changes of copy number. For these genes, the numbers showing inverse *trans* effect (increasing copy number with down-regulation or decreasing copy number with up-regulation) and direct *trans* effect (increasing copy number with up-regulation or decreasing copy number with down-regulation) **(Birchler and Veitia, 2012)** were balanced although bias was seen in several HE regions (**Figure 1d**; binomial test, p-value< 0.05). Gene ontology (GO) analysis showed that the two gene sets from HE-containing individuals showed distinctly enriched terms: while direct *trans* genes were associated with oxidative stress, peroxidase activity and kinase activity, inverse *trans* genes were related to molecular functions such as monooxygenase and hydrolase activity (**Figure S3**).

Genes located within the same chromosome regions but with variable copy numbers among different HE-containing individuals provided an opportunity to investigate whether there exists dosage compensation in gene expression. Four sets of candidate chromosome regions, including Chr3A (2 copies in the control plant, 3 copies in individual G2 and 4 copies in individual H), Chr3D (1 copy in individual G2, 2 copies in the control plant and 4 copies in individual H), Chr6A (2 copies in the control plant, 3 copies in individual J and 4 copies in individual I), Chr3D (1 copy in individual J, 2 copies in the control plant and 4 copies in individual I), were extracted to explore dosage compensation effect (**Figure S4a**). After strict statistics filtering (see Materials and Methods), we found that 9-174 (3.8-13.7%) genes showed constant expression levels among different HE-containing individuals with variable copy numbers. Also, the two subgenomes showed unequal propensities for dosage compensation, with subgenome A containing higher proportion of dosage compensating genes than subgenome D (**Figure S4b**; 11.8-13.7% versus 3.8-5.5%; Fihser’s exact test, p-value = 1.548e-11). Together, these results suggest that HEs impacted expression of HE-related gene primarily via *cis*-acting dosage effect, but *trans*-regulation also appeared to play an antagonistic role on some of these genes to maintain their constant expression levels, i.e., dosage compensation.

### Homoeolog expression bias contributes to expression variation after HE

We further explored how homoeolog expression bias (HEB) would influence changes of homeolog expression level and total expression level of homoeolog pairs in HE regions (**Figure 2a**). For clarity, only homoeologs with four copies from the same subgenome (two A copies were replaced by two D copies, and vice versa) in HE-containing individuals were used for further analysis. Based on the HEB patterns in the control plant (namely, no bias, A-bias and D-bias), homoeologs were divided into three classes: (*i*) no-bias, (*ii*) higher-homoeolog bias (higher-expressing homoeolog replaced the lower one) and (*iii*) lower-homoeolog bias (lower-expressing homoeolog replaced the higher one). We found that the proportion of dosage-insensitive homoeologs varied among the homoeolog classes (**Figure 2b**): subgenome A possessed the highest proportion of dosage-insensitive homoeologs in class *ii* compared with the rest two classes (on average 10.7%, 24.2% and 12.3% for class *i, ii* and *iii*, respectively; paired t-test, p-value < 0.05), whereas for subgenome D the proportion of dosage-insensitive homoeologs in class *ii* was also higher than class *i* but not than class *iii* (on average 10.3%, 19.4% and 16.7% for class *i, ii* and *iii*, respectively; paired t-test, p-value < 0.05). This observation suggests that homoelog members with equal contribution (no bias) are more likely dosage-sensitive, whereas the higher-expressing homoeologs are least subjected to dosage effect, which might be attributed to the impact of *trans*-regulation.

**Figure 2.**
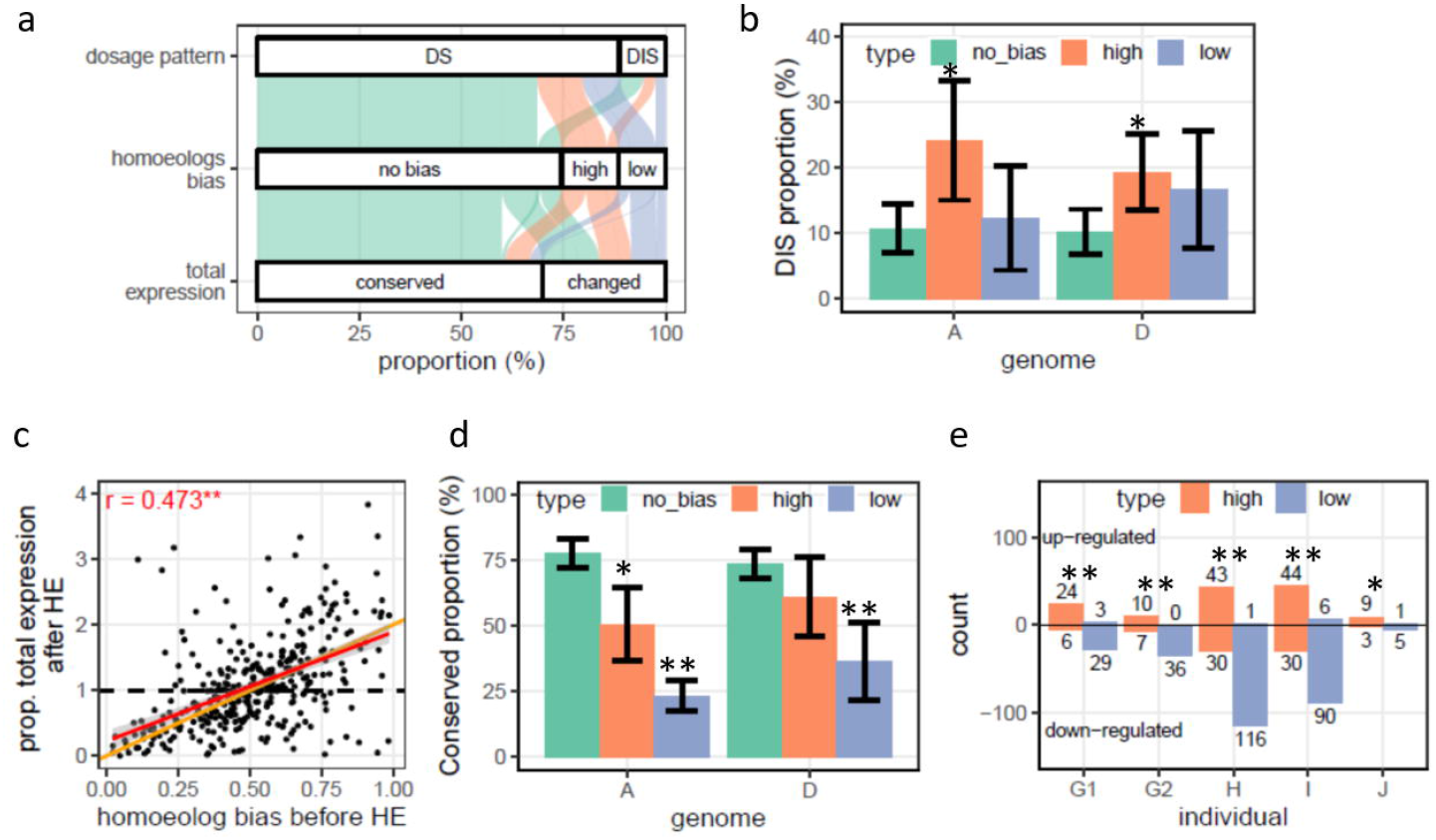
Expression variation of homoeolog pairs in HE regions. **(a)** The illustration that the effects of homoeolog expression bias (HEB, before HE) on DS and DIS genes and on total expression of homoeolog pairs (conserved and differentially expressed) after HE. **(b)** The DIS proportion in higher-expressing (denote as “high”; orange), lower-expressing (denote as “low”; blue) and no bias (green) homoeolog member sustained after HE in subgenomes A and D. Error bars indicate the variation of the proportion among HE-containing individuals (n = 5). The asterisks indicate a significant difference between no bias and “high” or “low” homoeologs (Paired t-test; *: p-value < 0.05). **(c)** The overall expression level of homoeolog pairs (after HE) versus HEB (before HE). The x-axis represents the degree of HEB and the y-axis represents the change ratio of total expression level. The black dashed line indicates the theoretical dosage-compensate pattern of total expression (y = 1). The orange line indicates the theoretical dosage-sensitive pattern of total expression (y = x). The asterisk indicates a significant correlation (Pearson’s product-moment correlation, p-value < 0.01). The red line fits the real data. **(d)** The conserved proportion in no bias, “high” and “low” homoeologs in subgenomes A and D. Error bars indicate the variation of the proportion among HE-containing individuals (n = 5). The asterisks indicate significant difference between no bias and “high” or “low” genes (Paired t-test; *: p-value < 0.05; **: p-value < 0.01). **(e)** Total expression changes in “high” and “low” homoeologs after HE. For each HE-containing individual, “high” and “low” homoeologs were assigned to totally up-regulated (above the x-axis) and down-regulated (below the x-axis) blocks, respectively. Digitals near each bar represent the gene count belonged to each block. The asterisks indicate significantly different enrichment between “high” and “low” homoeologs in totally up-regulated and down-regulated blocks (Fisher’s exact test; *: p-value < 0.05; **: p-value < 0.01).

We then investigated how HE affected total expression level of homoeolog pairs, i.e., whether the expression level of deleted homoeolog copies could be compensated for duplicated homoeolog copies. After HE, higher-expressing homoeolog tended to have higher total expression level of the pair (**Figure 2c and Figure S5**; r from 0.27 to 0.47; Pearson’s product-moment correlation, p-value < 0.01), which was mainly due to dosage effect. We therefore focused on how dosage effect affected the total expression level of the three classes of homoelog pairs. There was *ca*. three quarters of class *i* homoeologs maintaining original total expression level and were insensitive to HE (**Figure 2d**; 70.0-81.9% and 67.3-80.6% for subgenomes A and D, respectively). However, the proportion decreased in class *ii* homoeologs (37.1-64.8% and 45.0-85.7% for subgenomes A and D, respectively) and plunged to a much lower proportion in class *iii* homoeologs (15.4-27.8% and 26.9-62.5% for subgenomes A and D, respectively), reflecting lower-expressing homoeologs are more unlikely to maintain the original overall expression level (**Figure 2d**; paired t-test, p-value < 0.05). Moreover, for classes *ii* and *iii* homoeologs whose overall expression was changed, we observed that the higher-expressing homoeologs tended to show up-regulation whereas the lower-expressing homeologs tended to be further down-regulated (**Figure 2e**; Fisher’s exact test, p-value < 0.05 for all HE-containing individuals). These results suggest that HE leads to novel variation in overall expression level of homoeolog pairs due to dosage effect when HEB exists.

### Expression pattern of genes located in non-HE regions

All differentially expressed genes (DEGs) in non-HE regions between HE-containing individuals and the control were analyzed for *trans*-effects. In total, we identified 2,462 - 5,405 (10.3-21.7%) DEGs in response to HEs, in which subgenome D (9.6-20.3% DEGs) showed slightly higher degree of response than did subgenome A (10.9-23.0% DEGs) (**Table S1**; Fisher’s exact test, p-value < 0.001). This subgenome-asymmetric transcriptomic responses were associated with the phenomenon that more genes from subgenome D were replaced by subgenome A, i.e., higher levels of gene deletions (in subgenome D) might have stronger *trans*-effect on genes from the same subgenome (**Figure S6**). Meanwhile, more DEGs trended to be down-regulated (5.7-12.9%) than up-regulated (4.1% - 8.8%) in response to HEs (**Table S2**; binomial test, p-value < 0.001). GO enrichment analysis showed that down-regulated genes in all HE-containing individuals were enriched in GO terms such as cellulose biosynthetic and carbohydrate metabolic process (BP), hydrolase, heme, iron ion, oxidoreductase and peroxidase activity (MF), whereas up-regulated genes were enriched in variable GO terms among different individuals (**Figure S7**).

We next checked whether HE regions could also generate *cis*-effect on genes located on non-HE regions in the same recombined chromosomes. First, DEGs were divided into two classes based on their chromosomal locations: (*i*) DEGs on chromosomes including HE and (*ii*) DEGs on chromosomes without HE. We found that there is no significant difference in proportion between the two classes of DEGs (**Figure S8**; Fisher’s exact test and FDR correction, q-value > 0.05; except individual J). Second, we tested whether the length of HE was correlated with the proportion of DEGs among different chromosomes, and we found longer HE did not increase DEG proportion (**Figure S9**; Pearson’s product-moment correlation, p-value > 0.05; except individual G2). A previous study showed that chromosomes in polyploid wheat are unequal in their responses to aneuploidy with respect to the number of genes being dysregulated, and which was attributed to variation of *trans*-effect **(Zhang *et al*., 2017)**. Therefore, we tested whether such phenomenon also applies to the effect of HEs, which might be attributed to *cis*-effect. As shown in **Figure S10**, all chromosomes (including both recombined and non-recombined chromosomes) showed similar responses to HE except for chromosome 4D in individual I (Fisher’s exact test and FDR correction, q-value > 0.05), suggesting *trans*-effect acted almost equally on different chromosomes in HE-containing individuals. Together, these results suggest that positions of HE (*cis*-effect) play a minor role on the genes located in the same chromosome (excluding those located within HE-regions), reflecting gene expression variations on non-HE regions was mainly due to *trans*-effect or the combination of *trans*- and *cis*-effects.

### Homoeologs trended to co-respond to HE due to *trans*-regulation

The foregoing analyses of gene expression changes in non-HE chromosome regions were based on whole transcriptome which did not consider the possible differential responses between the subgenomes A and D. We next investigated whether the two subgenomes showed convergent or divergent regulation or subgenome-specific regulation in response to HE. To answer this question, 7,935 homoeolog pairs between subgenomes A and D were divided into five different classes based on their expression changes after HE: (*i*) both homoeolog members are DEGs that responded to HE in the same direction (convergent); (*ii*) both homoeolog members are DEGs but they responded to HE in opposite directions (divergent); (*iii*) only the homoeolog member of subgenome A is DEG (A-specific); (*iv*) only homoeolog member of subgenome D is DEG (D-specific); and (*v*) both homoeolog members are non-DEGs. We tabulated the five classes of homoeologs and made the following major observations. First, 10.6-23.0% (average 16.1%) of homoeolog pairs contained at least one DEG (classes *i* - *iv*, defined as HE-responsive homoeologs) in HE-containing individuals (**Figure 3a**). Second, most homoeolog pairs belonged to class *i* (39.0-46.0%), whereas <1% belonged to class *ii*. Notably, the proportion of class *i* homoeolog pairs was significantly higher than expected, suggesting HE-induced *trans*-effect plays a major role in co-regulating homoeologs expression. Third, 54.0-61.0% homoeolog pairs belonged to classes *iii* and *iv*, in which homoeolog members from subgenome D (24.1-27.8%) were more sensitive to HE than those from subgenome A (29.3-30.8%) except for individual G2 in which the reverse is observed (A:D = 32.2%:28.8%). Finally, after quantifying expression changes in different homoeolog classes, we found that class *i* showed more dramatic expression changes than classes *iii* and *iv* (**Figure 3b and Figure S11**; Mann-whitney-U test, p-value < 0.01).

**Figure 3.**
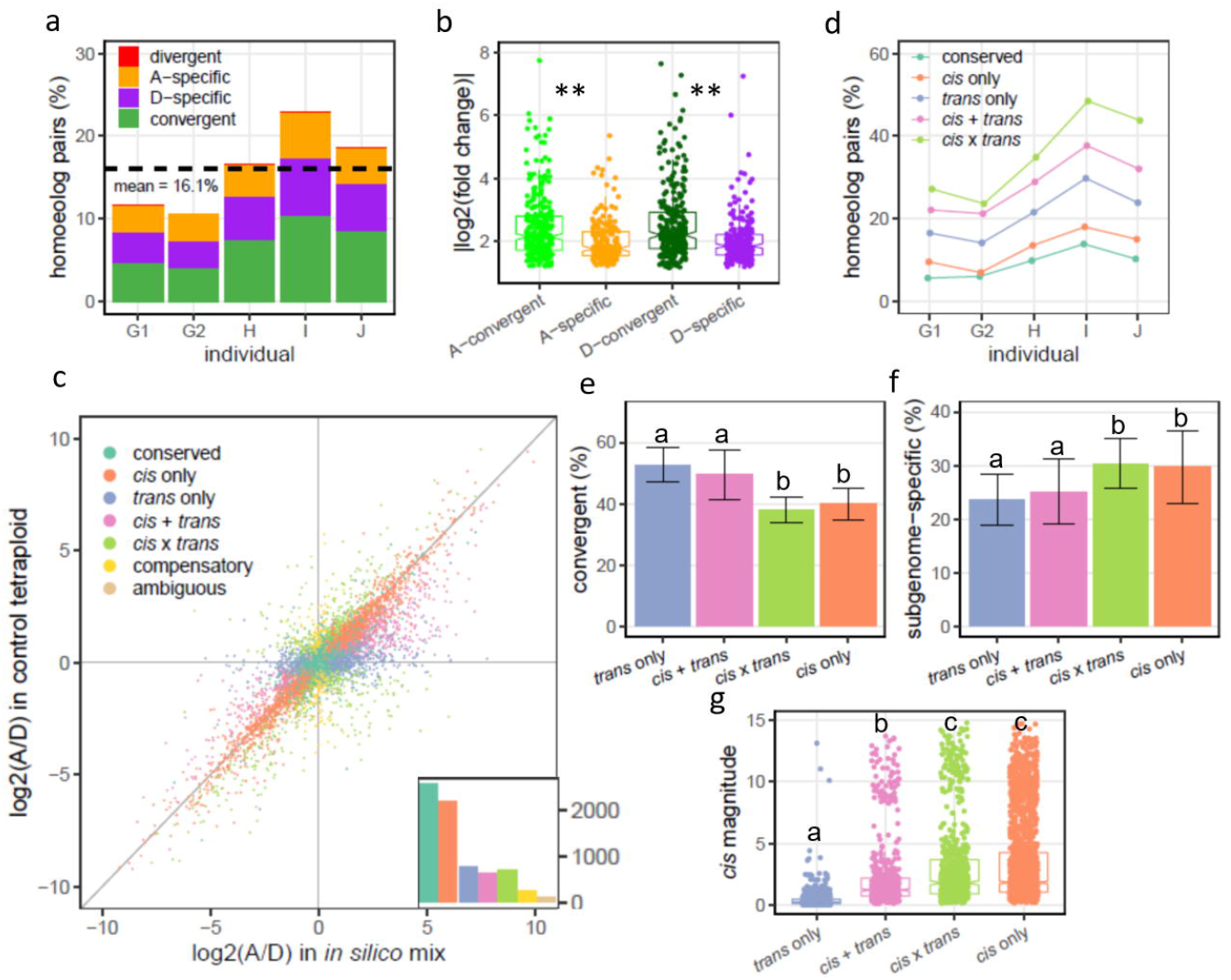
*cis/trans* regulation of homoeologs in non-HE regions. **(a)** Four expression classes of homoeolog pairs respond to HE. For each HE-containing individual, the ratio of homoeolog pairs in four different response classes including convergent (red), divergent (green), A-specific (orange) and D-specific (purple), were tabulated. The dashed line indicates the average response ratio among HE individuals. **(b)** Comparison of expression divergence between homoeologs in convergent and subgenome-specific classes of both subgenomes A and D, respectively, in individual G1. Y-axis represents the logarithm-transformed absolute value of expression ratio between G1 and control individuals. The asterisks represent a significant difference in expression ratio between convergent and subgenome-specific homoeologs in A and D subgenomes, respectively (Mann-Whitney-U test, p-value < 0.01). See other HE-containing individuals in **Figure S11**. **(c)** Homoeolog-specific expression (HSE) pattern between parents and control tetraploid. Logarithm-transformed expression ratio of *in silico* mix (the x-axis) versus control tetraploid (the y-axis) for 7953 homoeolog pairs is drawn. Seven HSE groups are shown with different colors. Homoeolog numbers falling into different groups are shown as a bar plot at the right bottom. **(d)** Homoeologs of five major HSE groups response to HE. **(e)** The proportion of convergent response and **(f)** Subgenome-specific response among four *cis*- and *trans*-related HEB groups. Error bars were calculated through five HE-containing individuals. The alphabets represent different levels among HSE groups (Paired t-test; **: p-value < 0.05). **(g)** *Cis* magnitude of homoeolog pairs among four *cis*- and *trans*-related HEB groups in control tetraploid. The alphabets represent different levels among HSE groups (Tukey and Kramer test after Kruskal-Wallis rank sum test, p-value < 0. 01).

We further asked whether and how original homoeolog-specific expression (HSE) pattern impacting response to HE, by measuring the expression pattern of homoeolog pairs in both *in silico* parental mix (reflecting the expression state in the A- and D-genome diploid parental species, *T. urartu* and *Ae. tauschii*) and the control plant (represents the expression state in AADD tetraploid) to distinguish seven types of homoeolog-specific expression (HSE) as defined **(Hu and Wendel, 2019)** (see Materials and Methods). In **Figure 4c**, homoeolog pairs related to conserved effect possessed the highest proportion (2,569, 35.4%), while 2,191 (30.2%) and 776 (10.7%) were associated with *cis* only and *trans* only effects, respectively. The remaining homoeolog pairs are 627 (8.6%) of *cis* + *trans*, 696 (9.6%) of *cis* x *trans* and 268 (3.7%) of compensatory effects, leaving 124 (1.7%) as unclassified (ambiguous). The following results were obtained. First, genes from different HSE patterns differed in their sensitivity to HE (**Figure 3d**), namely, homoeolog pairs related to *trans* effects (14.2% - 29.8% for *trans* only, 21.2% - 37.7% for *cis* + *trans* and 23.6% - 48.4% for *cis* x *trans* effects) are more sensitive in response to HE when compared with conserved (5.7% - 13.9%) and *cis* only effects (7.1% - 18.1%). Second, class *i* homoeolog pairs (convergent response to HE) were mainly enriched in *trans* only and *cis* + *trans* effects rather than *cis* only and *cis* x *trans* effects, whereas class *iii* and *iv* genes (subgenome-specific response to HE) showed an opposite trend (**Figure 3e-f**). This result is consistent with ranking of their original *cis*-effect magnitude (**Figure 3g**; Tukey and Kramer test after Kruskal-Wallis rank sum test, p-value < 0.01), that is, homoeolog pairs with lower *cis* magnitude (*trans* only effect) tended to show co-response to HE while those with higher *cis* magnitude (*cis* only effect) tended to manifest subgenome-specific response pattern.

**Figure 4.**
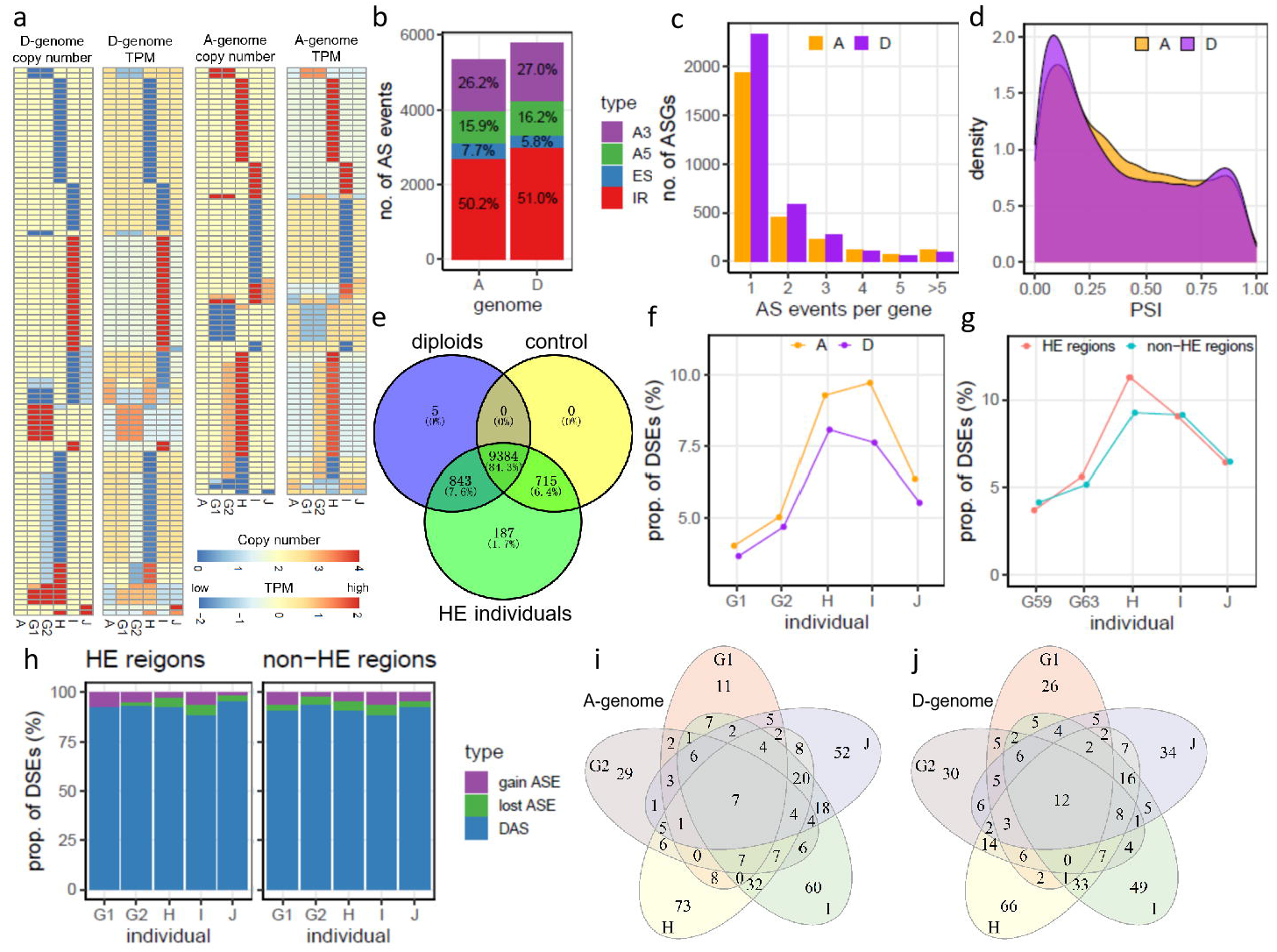
Characteristics of AS and their response to HE. **(a)** Copy number variation and corresponding expression changes of splicing-factor-encoding genes of A and D subgenomes. **(b)** The proportion of four types of AS events including alternative donor (AltA), alternative acceptor (AltD), exon skipping (ES) and intron retention (IR). **(c)** Distribution of ASGs associated with the different number of AS events. **(d)** PSI distribution between subgenomes A and D. **(e)** Venn diagram of AS events among in *silico mix*, control tetraploid and HE-containing individuals. **(f)** and **(g)** The proportion of DSEs **(f)** between subgenomes and **(g)** between HE and non-HE regions. **(h)** The proportion of qualitative DAS (including gain AS event and lost AS event) and quantitative DAS in HE and non-HE regions. **(i) and (j)** Venn diagram of DAS among HE-containing individuals in subgenomes **(i)** A and **(j)** D.

### Global profiling of alternative splicing events related to HE

Alternative splicing (AS) plays important roles in producing different transcripts from the same gene and thereby generating transcriptome diversity. Our aforementioned results show that HE caused expression changes of large number of genes; of which 195 splicing-related genes, 81 and 104 from subgenomes A and D, respectively, were identified (**Figure 4a**). This prompted us to further analyze the effects of HE on the regulation of global AS patterns.

We profiled AS events using the same transcriptome data as for gene expression analyses, described above. Based on a series of bioinformatics processes with stringent criteria (Materials and Methods), 11,134 AS events were identified from 6,367 AS genes (defined as ASGs). There are 5,346 (from 2,924 ASGs) and 5,788 (from 3,443 ASGs) AS events derived from subgenomes A and D, respectively (**Figure 4b**). We found compositions of AS types were similar between the two subgenomes (**Figure 4b and Table S3**): IR (intron retention) was the most common AS event (50.2% - 51.0%), followed by A3 (alternative acceptor; 26.2 – 27.0%), then A5 (alternative donor; 15.9% - 16.2%) and the least was ES (exon skipping; 5.8% - 7.7%). The number of AS events per ASG showed nearly identical distribution between subgenomes A and D (**Figure 4c**; Kolmogorov-Smirnov test, p-value = 0.931). Majority of ASGs contained only one AS event with a slight dominant proportion in subgenome D (66.4% vs. 67.7% AS genes from subgenomes A and D, respectively), whereas only 4.2% and 2.8% ASGs related to multiple AS events (>5 AS events per gene) in subgenomes A and D, respectively, which are significantly different (Fisher’s exact test, p-value = 3.56e-3). When quantifying the degree of AS events, the distribution of PSI (percentage of splicing index) showed significant difference between the two subgenomes (Kolmogorov-Smirnov test, p-value <2.2e-16), in which subgenome A possessed more AS events with medium level (0.25 < PSI < 0.75) whereas more AS events with extreme level (PSI < 0.25 or PSI > 0.75) were identified in subgenome D (**Figure 4d**). These results suggest that the two subgenomes in the AADD tetraploid plants possessed analogous AS atlas with subtle variations.

We previously found in a rice synthetic segmental allotetraploid that rampant HEs contributed to large-scale homozygosity of polyploid genome, in which extensive variation of AS, and hence transcriptome diversity, occurred **(Zhang *et al*., 2019)**. To test whether such phenomenon also occurred in the HE-containing AADD tetraploid wheat and to what extent HE contributed to transcriptome diversity, a qualitative analysis was performed by comparing the numbers of AS events among diploid parents, control tetraploid and the variable HE-containing tetraploid individuals. As shown in **Figure 4e**, it is clear that (*i*) 84.3% (9,384 out of 11,134) AS events (with PSI > 0.05) were overlapped among three groups; (*ii*) there are rare diploid-specific or tetraploid-specific AS events, whereas HE contributed to ~1.7% (187 out of 11,134 AS events) transcriptome diversity. Quantitative analysis also demonstrated that AS signal is relatively stable in A3, A5 and ES but with slight changes in IR, after HE (**Figure S12**; Kruskal-Wallis-post-hoc-test, p-value < 0.05).

Similar to gene expression, we further investigated the characteristics of AS response to HE by comparing AS variations between control and variable HE-containing individuals. In total, 408 - 819 (4.1-9.6%) DSEs (differentially splicing events, ΔPSI > 0.2 or ΔPSI < −0.2; see Materials and Methods) were identified, in which the proportion of subgenome A was higher than that of subgenome D (Fisher’s exact test, p-value < 0.05 for individual H, I and J; **Figure 4f**). Intriguingly, the proportion of DSEs in individual H and I were higher than other individuals, which should be attributable to their relatively more numerous HEs. To test whether DSEs are sensitive to genomic locations, we compared the proportions of DSEs between HE (*cis* regulation) and non-HE (*trans* regulation) regions and observed no discernible difference (**Figure 4g**; Fisher’s exact test, p-value = 0.024 for individual H; p-value > 0.05 for another four individuals). Not surprisingly, majority of the AS events belonged to quantitative changes (**Figure 4h**; 88.7-96.3% and 90.7-95.8% for HE and non-HE regions, respectively) rather than the qualitative DSEs (i.e., gain vs. loss of AS events, which were 3.7-11.3% and 4.2-9.3% for HE and non-HE regions, respectively). This is in stark contrast to polyploidy- or domestication-induced AS variance in natural polyploid wheat species **(Yu *et al*., 2020)**. We then focused on the conservation of AS changes in response to HE by comparing conserved DSEs (on non-HE regions) among five HE-containing individuals. As shown in **Figure 4i**, only 1.8% (7 out of 391, subgenome A) and 3.3% (12 out of 368, subgenome D) DSEs were shared among all individuals, whereas 57.5% (225 out of 391, subgenome A) and 55.7% (205 out of 368, subgenome D) DSEs were individual-specific, suggesting the HE-induced copy number variation of variable splicing factors might generate distinct *trans*-effects and act on different gene loci among different individuals. Further RT-PCR experiment also accurately validated the manners of HE-induced DSEs (**Figure S13**).

### Homoeo-AS events in response to HE

We next explored how homoeo-AS events (AS events identified from conserved exon-intron junctions of homoeolog mates; see Materials and Methods) respond to HE. In 2,443 homoeolog pairs (4,886 genes) from 6,367 ASGs, nearly three-quarters (1,808) homoeolog pairs lacked corresponding AS events (**Figure 5A**), suggesting intrinsic AS partitioning between subgenomes occurred in the synthetic tetraploid wheat, which might be attributed to the high degree of divergence between parental species, *T. urartu* and *Ae. tauschii*. We next mainly focused on the remaining 635 homoeolog pairs possessing 969 high confidence homoeo-AS events, including 381 IR, 346 A3, 213 A5 and 29 ES event pairs, to explore their regulation fatal after HE.

**Figure 5.**
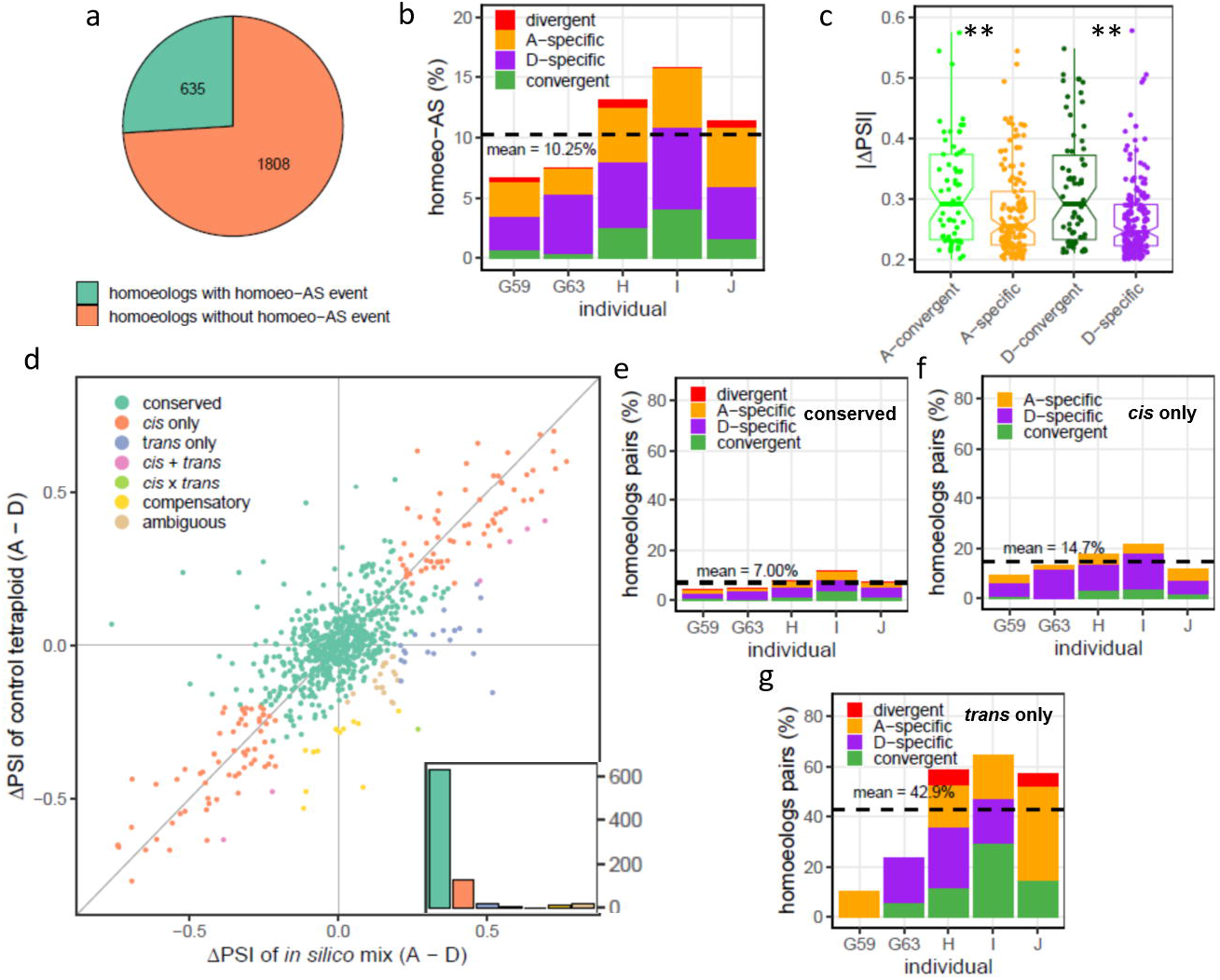
Homoeo-AS event response to HE. **(a)** The proportion of homoeologs with and without homoeo-AS event. **(b)** Four classes of homoeo-AS events response to HE. The dashed line indicates the average response ratio among HE-containing individuals. **(c)** Comparison of AS divergence between homoeo-AS events in convergent and subgenome-specific classes of both subgenomes A and D, respectively. Y-axis represents the absolute Δ PSI value of AS ratio between HE-containing individuals and control tetraploid. The asterisks represent a significant difference in AS ratio between convergent and subgenome-specific homoeo-AS events in subgenomes A and D, respectively (Mann-Whitney-U test, p-value < 0.05). **(d)** Homoeolog-specific splicing (HSS) pattern between parents and control tetraploid. absolute ΔPSI value of *in silico* mix (x-axis) versus control tetraploid (y-axis) for 635 homoeo-ASEs is drawn. Seven HSS groups are shown with different colors. The number of homoeo-AS events falling into different groups is shown as a bar plot at the right bottom. **(e)**, **(f)** and **(g)** Homoeologs of three major HSS groups response to HE, including **(e)** conserved, **(f)** *cis* only and **(g)** *trans* only effects. The dashed line indicates the average response ratio among HE-containing individuals.

We asked whether homoeo-AS events showed convergent response to HE. As shown in **Figure 5b**, HE-induced DSEs in homoeo-AS events (6.6-15.1%, average 10.3%) was much less responsive to HE than DEGs in homoelog pairs (10.6-23.0%, average 16.1%). Unlike homoeolog expression changes, DSEs mainly manifested as subgenome-specific response pattern (47-82 representing 76.292.7% homoeo-AS events), including 16 - 40 (29.1-44.4%) A-specific and 22 - 47 (36.7-63.6%) D-specific homoeo-AS events, rather than convergent response pattern (5.5-25.9%) (**Figure 5b**). Compared with other individuals, individual H and I possessed significantly higher proportions of variable homoeologous AS pairs, corresponding with the proportion of all DAS events. Notably, similar to homoeolog expression changes, convergent homoeo-AS events showed greater response (measured as ΔPSI) than subgenome-specific AS events (Mann-Whitney-U test; p-value = 0.01 for subgenome A and p-value = 3.47e-4 for subgenome D), but there is no obvious difference between the two subgenomes (**Figure 5c**).

Then, we analyzed whether HE-induced DSE in homoeo-AS is associated with *cis* and/or *trans* regulation. First, similar to the definition of HSE, *cis* and/or *trans* regulation of homoeo-AS events (namely homoeolog-specific splicing, HSS) was classified into seven regulatory categories, based on the divergence of each homoeo-AS event in both *in silico* mix and the control tetraploid (see Materials and Methods). We found that (**Figure 5d**): (*i*) the overall distribution of splicing difference within homoeo-AS events in the control tetraploid was significantly correlated with that in the *in silico* mix (Pearson’s product-moment correlation test, p-value < 0.001, r = 0.78); (*ii*) except for the majority of conserved homoeo-AS events (76.5%, 627 out of 820), the number of *cis* only homoeo-AS events (15.9%) is overwhelmingly greater than the rest categories (totally 7.6%). Second, we checked how homoeo-AS events from different *cis/trans* regulation categories in response to HE, especially in conserved, *cis* only and *trans* only categories. Compared with conserved category, homoeo-AS events belonging to the *cis* only category was more responsive to HE (**Figure 5e**; average 14.7% vs. 6.0% DSEs), in which D-specific homoeo-AS events contributed more than the rest homoeo-AS patterns, inclduing A-specific and convergent). Notably, homoeo-AS events belonging to the *trans* only category exhibited the highest level of response to HE (average 42.9% DSEs), although different individuals possessed variable proportions of responsive patterns (**Figure 5e**). These results suggest that although *cis*-regulation and/or parent legacy play major roles in regulating HSS at the early stages of polyploidization, homoeo-AS events with *trans*-regulation are more susceptible to allele copy number variation and expression changes of *trans* splicing factors introduced by HE.

## Discussion

HE provides the material basis for rapid karyotype variation, which bears profound physiological and phenotypic consequences, and relevant to adaptive evolution in allopolyploid species **(Hollister, 2015, Gou *et al*., 2018, Wu *et al*., 2021)**. Nevertheless, the direct genetic effects of HE on genome and transcriptome remain to be systematically studied. In this respect, synthetic allopolyploids provide suitable systems because additional evolutionary forces, selective or neutral, are yet to happen. In this study, we focused on how RNA-transcription response to rampant HE via *cis* and *trans* effects in a synthetic allotetraploid wheat (genome AADD), based on our previous work reporting on the mechanism and features of HE generation **(Zhang *et al*., 2020)**.

### Effect of HE on gene expression in *cis*

Gene copy number variation (CNV) may cause either dosage effect or dosage compensation on gene expression **(Guo *et al*., 1996, Acar *et al*., 2010, Birchler and Veitia, 2012, Malone *et al*., 2012, Zhang *et al*., 2017, Hou *et al*., 2018)**. In allopolyploids, HE also leads to local CNV in *cis* homoeologous genomic regions **(He *et al*., 2017, Stein *et al*., 2017, Hurgobin *et al*., 2018, Wu *et al*., 2021)**. Thus the issue arises with respect to whether HE-induced substitution of homoeolog pairs compensates for their original total expression level and maintains transcriptome homeostasis.

Our results showed that dosage effect plays a major role on gene expression within HE regions, leading to variable overall expression of homoeolog pairs after HE when homoeolog expression bias (HEB) presents. Similar phenomenon was found in *Brassica napus* **(Lloyd *et al*., 2018)**, suggesting HE-induced *cis* genetic consequences are conserved in different synthetic and natural allopolyploid plant species. Considering the high proportion of no-bias homoeologs, changes in their overall expression level via HE seems to be limited. Therefore, it can be envisioned that allopolyploids possessing high proportion of HEB homoeologs might seriously affect transcriptome stability when HE occurred. Notably, we found that some HEB homoeologs remained the same expression state in the HE-containing tetraploid individuals, suggesting *trans* effect of HE could also offset the transcriptome load incurred by the *cis*-effect of HE and maintain transcriptome homeostasis.

### HE-induced *trans*-regulation on gene expression

The duplication and/or deletion of whole chromosome(s) in aneuploidy can induce dysregulation of genes residing on non-homologous chromosomes, which is manifested as *trans* regulation **(Letourneau *et al*., 2014, Zhang *et al*., 2017, Hou *et al*., 2018, Zhu *et al*., 2018)**. For HE, it might change the gene regulatory machinery in allopolyploids because of asymmetric gene content among homoeologous chromosomes and sub-functionalization and new-functionalization of homoeologs **(Feldman *et al*., 2012, Panchy *et al*., 2016, Pont and Salse, 2017)**. Therefore, the combined effects of local copy number redundancy and gene loss in *trans* factors (such as transcription factors and splicing-related factors) could also trigger variations of *trans* genomic environment **(Hurgobin *et al*., 2018)**.

Our results suggest dosage imbalance of genes on HE segment could further induce rapid response at *trans* genomic regions, suggesting the behavior of homoeolog pairs response to HE is associated with their original *cis/trans* regulation pattern. Indeed, after merging divergent genomes in the same nucleus, expression divergence of alleles (hybrid) and/or homoeologs (allopolyploid) with conserved *cis* elements will be attenuated or eliminated by *trans* environment **(Mcmanus *et al*., Xu *et al*., 2014, Hu and Wendel, 2019)**. Therefore, when HE induces CNV of *trans* factors, it is expected that the target homoeolog pairs with *trans*-effect will show convergent expression response whereas those with divergent *cis* element might be subjected to subgenome-specific regulation.

### AS response to HE

AS was observed to respond to changes of external conditions (such as abiotic/biotic stresses) and internal genomic environment (including WGD and structure variations) in plants **(Staiger and Brown, 2013, Laloum *et al*., 2018, Gao *et al*., 2021)**, which contribute to diversity and complexity of functional genome through enlarging the variety of RNA isoforms **(Graveley, 2001, Syed *et al*., 2012)**. HE-induced AS response provides an extra dimension to further understand the genetic consequences of allopolyploidy.

Our results provided a solid evidence that HE-induced CNV of splicing factors could re-program their transcription atlas, providing a causal link between HE and AS. Notably, HE plays a role on fine-tuning the transcriptome diversity (about 2%), which however is to a less magnitude than we observed in a rice segmental allopolyploid **(Zhang *et al*., 2019)**. We noted however that the effect of HE on AS is milder than that of evolutionary forces such as natural and artificial selection (domestication) in wheat. For example, it was shown that domestication in wheat and other crops has led to dramatic AS changes including large-scale AS loss and *de-novo* generation of new transcript isoforms **(Yu *et al*., 2020)**, probably because HEs may underpin desired agronomic traits and hence are strongly selected for **(Zhao *et al*., 2006, Stein *et al*., 2017, Bertioli *et al*., 2019, Raman *et al*., 2021)**.By contrast, we found the effect of HE on AS is primarily quantitative. Notwithstanding, HE-imposed transcript diversity might play important roles at the critical initial stages of allopolyploid formation to enlarge the adaptive phenotypic space when the intrinsic genetic depauperation may severely preclude establishment of newly formed allopolyploid individuals.

In sum, our study shows that asymmetric homoeologous chromosomal segmental replacement mediated by HE causes rapid and extensive changes in gene expression and AS, due both to *cis* effects on genes located within HE regions also *trans* regulation of genes in non-HE regions of the genome, which together may contribute to adaptive evolution of newly formed allopolyploids.

## Materials and Methods

### Plant materials

The tetraploid wheat (AADD) was produced by intergeneric hybridization between *T. urartu* (A) and *Ae. tauschii* (D) followed by colchicine-mediated genome doubling. A single S2 euploid individual seed was used as the founder to construct the S12 tetraploid wheat population (**Gou *et al*., 2018**). Parent diploids, control tetraploid in S6 generation and five HE-containing tetraploids from S9 generation, were sampled from the original population **(Zhang *et al*., 2020)**. All individual plants were successfully karyotyped based on fluorescence *in situ* hybridization (FISH) and genomic *in situ* hybridization (GISH). All AADD plants and corresponding diploid parents (A and D) were grown in a glasshouse under normal conditions (25: 20°C, 16 : 8 h, day : night). Young leaves were immediately frozen in liquid nitrogen. Total RNA was isolated using Trizol (Invitrogen) based on standard protocol.

### Data collection and quality control

Library construction, cluster generation and Hiseq2500 sequencing (Illumina) were carried out with standard protocols for both DNA and RNA samples. Trimmomatic (Bolger, et al. 2014) was used to filter out adapters and low-quality reads (reads with Phred quality less than 10). Clean data have been deposited the SRA database (http://www.ncbi.nlm.nih.gov/sra/) with accession number PRJNA608801.

### Expression dosage analysis in HE-containing AADD tetraploids

After quality control, high-quality RNA-seq data were aligned to the AADD reference genome using STAR (v2.5.4) **(Dobin *et al*., 2013)** with two-round alignment method. Only sequencing reads that uniquely aligned were retained for further analysis. HTseq-count **(Anders *et al*., 2015)** was used to count reads number for each annotated gene model and counts matrix was constructed. TPM (<u>T</u>ranscripts <u>P</u>er <u>M</u>illion reads) values were used for measuring gene expression level. Genes with TPM > 1 in at least two experiments were used for further analysis.

To evaluate the contribution of dosage effect, HE-related genes with different copy number (including one copy, three and four copies) and with different subgenome origination (A and D) in each HE-containing individual were compared separately with those in control plant using DESeq2 **(Love *et al*., 2014)** as described in **(Lloyd *et al*., 2018)**. To directly compare genes with different copy number, the size factors were re-estimated. Briefly, genes located on none HE regions were used to estimate original size factor. Then, for HE-related genes, new size factors were generated by multiplying their corresponding DNA copy number by original size factors from genes in non-HE regions. The new size factors were used for further DESeq2 analysis. Genes with q-value <0.05 and fold change > 2 were defined as DEGs (differentially expressed genes). For each comparison, DEGs were treated as dosage-insensitive (DIS) genes, whereas the remaining genes were dosage-sensitive (DS) genes.

To identify genes associated with dosage compensation among genomic regions of different copy numbers among different HE-containing individuals, method from Zhang *et.al* were performed with several modification **(Zhang *et al*., 2017)**. First, Pearson’s product-moment correlation test was performed to test if copy number is positively correlated with expression level for each gene. Genes without significant correlation coefficient (p-value > 0.05) were retained for further filtering. Second, for each gene, one-way analysis of variance was performed to test if gene with difference copy numbers was significantly different expression. Genes with p-value larger than 5% were treated as dosage compensation genes. Using this method, we tested four genomic regions including Chr3A (2 copies in control plant, 3 copies in individual G2 and 4 copies in individual H), Chr3D (1 copy in individual G2, 2 copies in control plant and 4 copies in individual H), Chr6A (2 copies in control plant, 3 copies in individual J and 4 copies in individual I) and Chr3D (1 copy in individual J, 2 copies in control plant and 4 copies in individual I).

### Analysis of homoeolog pairs

Homoeolog pairs were identified by jcvi utility libraries (https://github.com/tanghaibao/jcvi) based on sequence similarity and genomic position of genes from A and D subgenomes. To identify homoeolog bias at expression level, Student’s t-test was performed to compare the TPM difference in each homoeologous gene pair. After FDR (false discovery rate, using Benjamini-Hochberg procedure) correction, homoeolog pairs with q-value < 0.05 were associated with biased expression (A-bias or D-bias). Student’s t-test was also performed to compare the difference between total expression of each homoeolog pair before and after HE. Total expression levels with q-value < 0.05 and fold change > 2 between control and HE-containing individuals were defined as differentially overall expression homoeolog pairs.

### *Cis*- and *trans*-regulation analysis in control tetraploid

As described in **(Hu and Wendel, 2019)**, homoeolog-specific expression (HSE) analysis was performed to classify homoeolog pairs with different regulation patterns. Briefly, expression difference of orthologs A and D in diploid parents are attributed to the combination of both *cis*- and *trans*-effects (*cis* + *trans*, marked as a). Meanwhile, in a common *trans* environment, homoeolog expression divergence in tetraploids are attributed to the pure *cis* effect (*cis*, marked as b). Therefore, the *trans* effect can be estimated from subtracting a from b (*trans*, marked as a - b). For each homoeologous gene pair, student’s t-test was performed to test the significance of *cis* + *trans* effects (a, TPM difference between orthologs A and D in *in silico* mix tetraploid), *cis* effect (b, TPM divergence between homoeologs A and D in control tetraploid) and *trans* effect (a-b, log2-transformed A:D ratio between *in silico* mix and control tetraploids), respectively. All three types of p-values were then corrected by FDR method with rate of 0.05. Totally 7,953 expressed homoeologous gene pairs were classified to six groups: (i) *cis* only: significant difference in b but not in a-b; (ii) *trans* only: significant difference in both a and a-b but not in b; (iii) *compensatory*, significant difference in both b and a-b but not in a; (iv) *cis* and *trans*, significant difference in a and b and a-b; (v) *conserved*: no significant difference in a and b and a-b and (vi) ambiguous: homoeolog pairs which don’t meet any of the above situations.

### Alternative splicing (AS) analysis

Transcript structure annotation file was reconstructed and filtered as follow steps. First, isoforms from each tetraploid replicate (including *in silico* Mix) were assembled separately (genome-guided mode) and then merged together using Scallops with the following parameters: --min_transcript_coverage 3 --min_flank_length 5 **(Pertea *et al*., 2015)**. Next, transcripts with each junction more than three supported reads (junction splicing reads) in each replicate samples were maintained according to splicing junction (SJ) information generated by STAR program. Then, Stringtie (Pertea *et al*., 2015) was performed to merge all isoforms from all replicate samples using the parameters: --merge -i -f 0.05 -T 1 -F 1 -m 200. Finally, Gffcompare (https://ccb.jhu.edu/software/stringtie/gffcompare.shtml) was used to compare the reconstructed annotation with the union of original genome annotation files (GFF file of *T. urartu* and *Ae. tauschii* genome from http://www.mbkbase.org/Tu/ and http://aegilops.wheat.ucdavis.edu/ATGSP/annotation/, respectively). Only isoforms with the class code of “=” and “j” were kept for further filtering step. Totally, 65,927 isoforms were obtained for further identification of AS events.

AS events were identified using SUPPA2 **(Trincado *et al*., 2018)** based on above annotation file. Four major types of AS event including intron retention (IR), exon skipping (ES), alternative donor (A5) and alternative acceptor (A3) and corresponding junction coordinates were acquired. To remove false-positive AS events, custom Python scripts were performed to count corresponding junction reads for each ASE in each replicate sample. For each AS event, sequencing reads related to inclusion and exclusion junctions were defined as inclusion and exclusion reads, respectively. AS events including at least three inclusion reads in at least one individual were kept, and inclusion and exclusion reads of three replicates from the same individual were added up, respectively. Finally, 11,135 high confident AS events were kept. Meanwhile, PSI (Percent Spliced Index) value which quantified the degree of AS event was also calculated. To obtain HE-induced DSE (Differential Splicing Event), binomial test was performed for each AS event based on inclusion and exclusion reads count between control and HE-containing individuals followed by FDR correction. DSE was defined if the q-value < 0.05 of corresponding AS events and the absolute value of ΔPSI > 0.2.

To identify homoeologous AS event pairs (homoeo-AS events) and quantify their divergence, continuous exons which associated with AS event in homoeolog genes were extracted and treated as target exon sets. For each target exon set, all exons from its homoeolog mate were treated as query sequence and aligned to it to identify homoeologous continuous exons pairs using BLASTn (-task dc-megablast). AS event which occurred in continuous exons pairs were defined as homoeo-AS event. Meanwhile, HSB (Homoeologs Splicing Bias) analysis was also performed according to binomial test between two mates of homoeo-AS event. HSB was defined when the model with p-value < 0.05 and the absolute value of Δ PSI > 0.2.

The homoeolog-specific splicing (HSS) analysis pipeline was constructed similar to homoeolog-specific expression (HSE). AS difference between ortholog AS events A and D in diploid parents is attributed to the combination of both *cis*- and *trans*-effects (*cis* + *trans*, marked as a). Meanwhile, in a common *trans* environment, divergence between two mates of homoeo-AS event in control tetraploid is attributed to the pure *cis* effect (*cis*, marked as b). The *trans* effect of each homoeo-AS events can be estimated from subtracting a from b ((*trans*, marked as a - b). For each homoeo-AS events, (1) binomial test was performed to check the significance of *cis* + *trans* effects (a, AS difference between ortholog AS events A and D in *in silico* mix tetraploid) and *cis* effect (b, HSB between homoeo-AS events A and D in control tetraploid); (2) the threshold of *trans* effect is set to 0.2 (namely a - b > 0.2). Totally 969 homoeo-AS events were classified to six groups: (*i*) *cis* only: significant difference in b but not in a-b; (*ii*) *trans* only: significant difference in both a and a-b but not in b; (*iii*) *compensatory*, significant difference in both b and a-b but not in A; (*iv*) *cis* and *trans*, significant difference in a and b and a-b; (*v*) *conserved*: no significant difference in a and b and a-b and (*vi*) ambiguous: homoeo-AS events which don’t meet any of the above situations.

RNA samples of all individuals were reverse-transcribed by Trnascript One-Step gDNA Removal and cDNA Synthesis SuperMix (TransGen) and used for RT-PCR validation of ASEs. Ten randomly ASEs (IR events) were chosen for further validation of ASEs between HE-containing individuals and control plant. Related primer sequenced were shown in **Table S4**.

## Acknowledgements

This study was supported by the National Natural Science Foundation of China (NSFC #32100179 to ZZ, NSFC #31830006 to BL and NSFC #31970238 to LG).

## Author Contributions

Z.B.Z., L.G., and B.L. designed the research. Z.B.Z., H.W.X., X.W.G., N.L., L.G. and B.L. performed the research. Z.B.Z., H.W.X., X.T.M., J.Z.L. and R.L.L. analyzed data, H.W.X. and J.Z. did experimental validation. Z.B.Z., L.G., and B.L. wrote the manuscript.

