## supplementary figures and tables for "Effects of homoeologous exchange on gene expression and alternative splicing in a newly formed allotetraploid wheat"

Figures S1 to S13

Tables S1 to S4

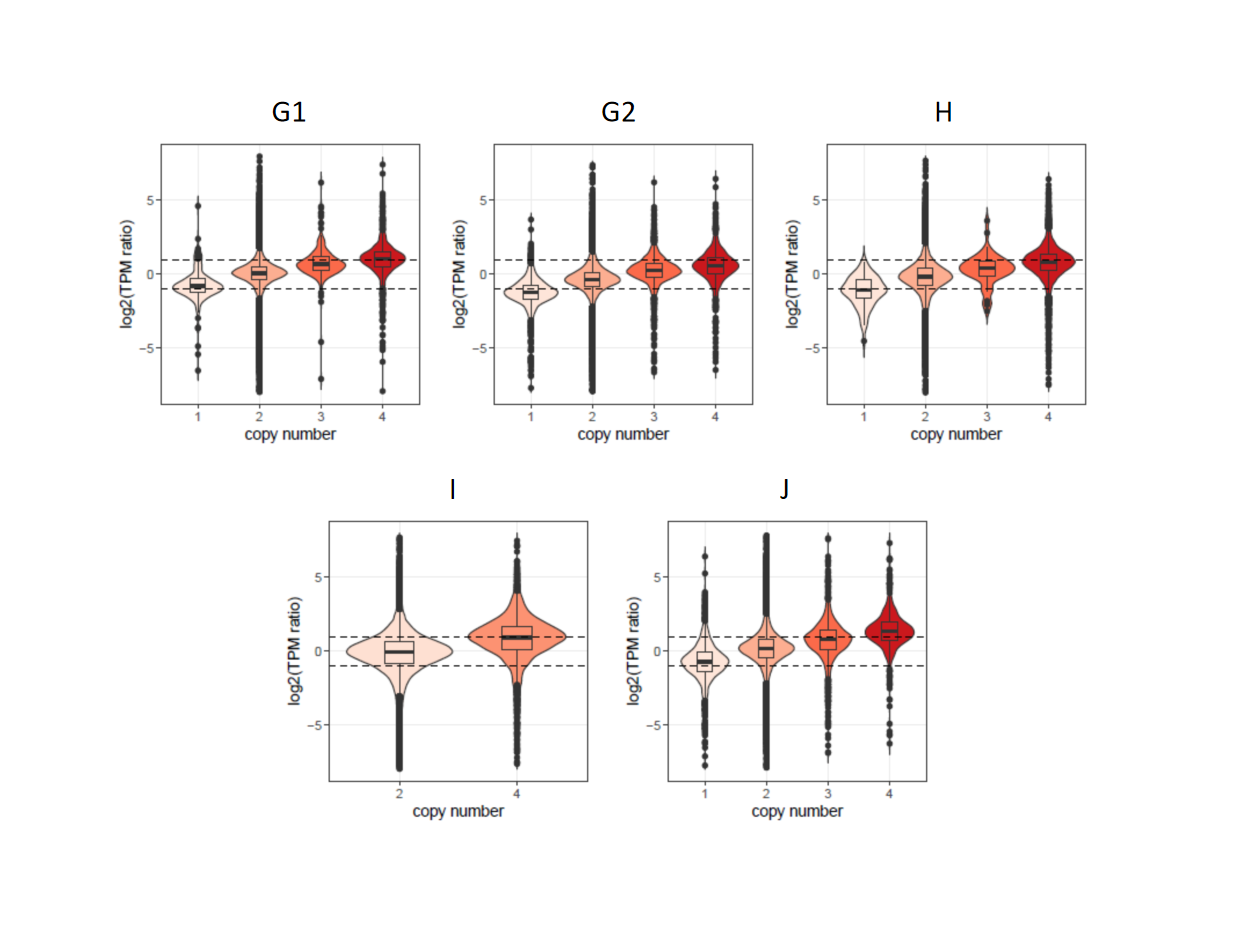

**Figure S1. Relationship between gene expression level and corresponding genomic copy number in five HE individuals.** X-axis represents gene copy number. Y-axis represents Log2-transformed expression fold changes between HE samples and control sample at different copy number levels. Dash lines represents fold changes of 2 and 0.5.

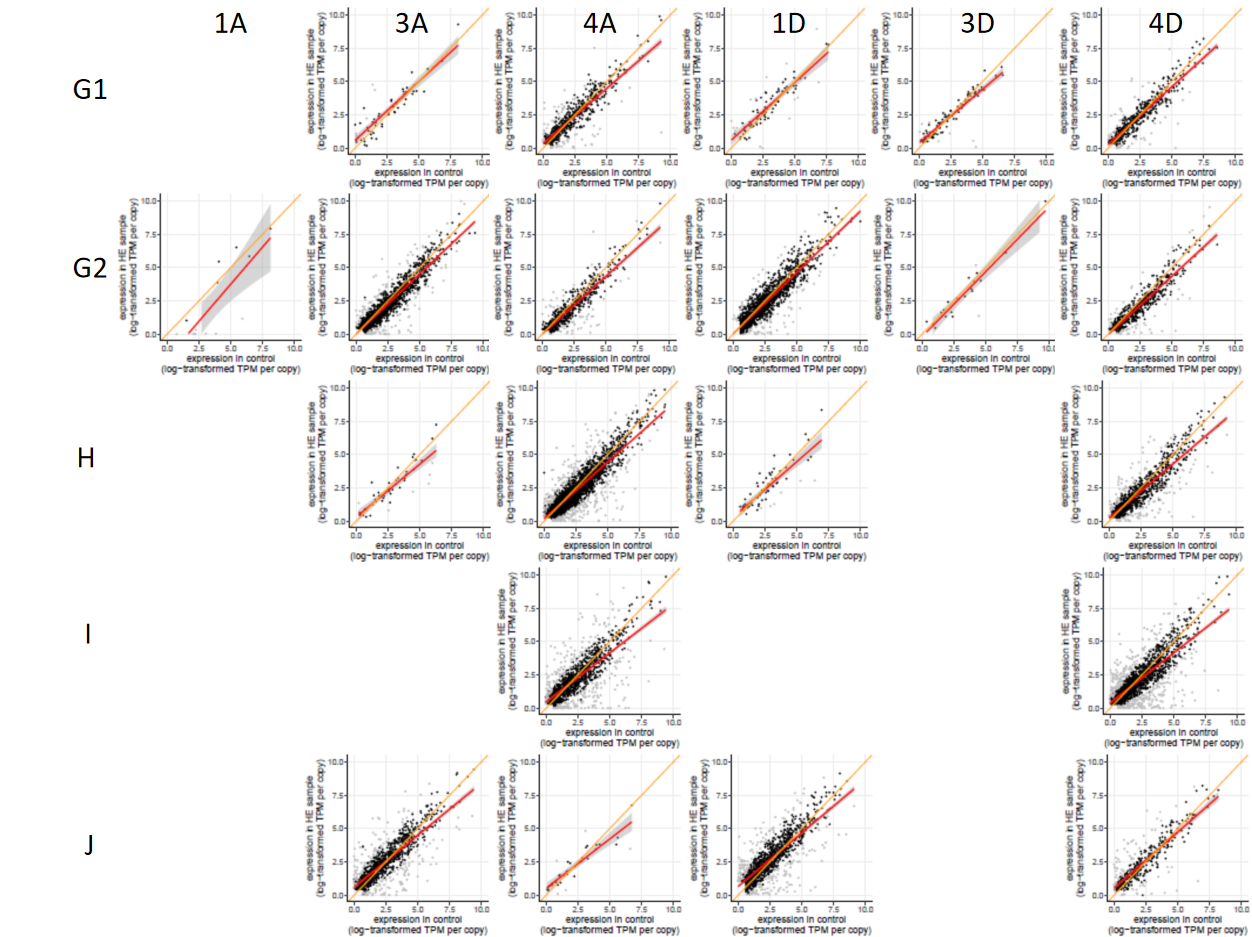

**Figure S2. Per copy expression level between HE samples and control sample at different copy number level.** For each panel, the x-axis represents the expression level divided by corresponding copy number (here is two) in control tetraploid, and the y-axis represents the expression level divided by corresponding copy number. Black and grey dots indicate genes with dosage-sensitive (DS) and non-dosage-sensitive (NDS) expression, respectively. Red line represents the fitting curve based on real data, whereas the orange line indicates the theoretical dosage-sensitive trendline (y = x). “1A” (“1D”) means one copy of gene belonged to AA (DD) subgenome was replaced by one copy from DD (AA) subgenome. “3A” (“3D”) and “4A” (“4D”) mean one copy and two copies of gene belonged to DD (AA) subgenome was replaced by one copy and two copies from AA (DD) subgenome, respectively.

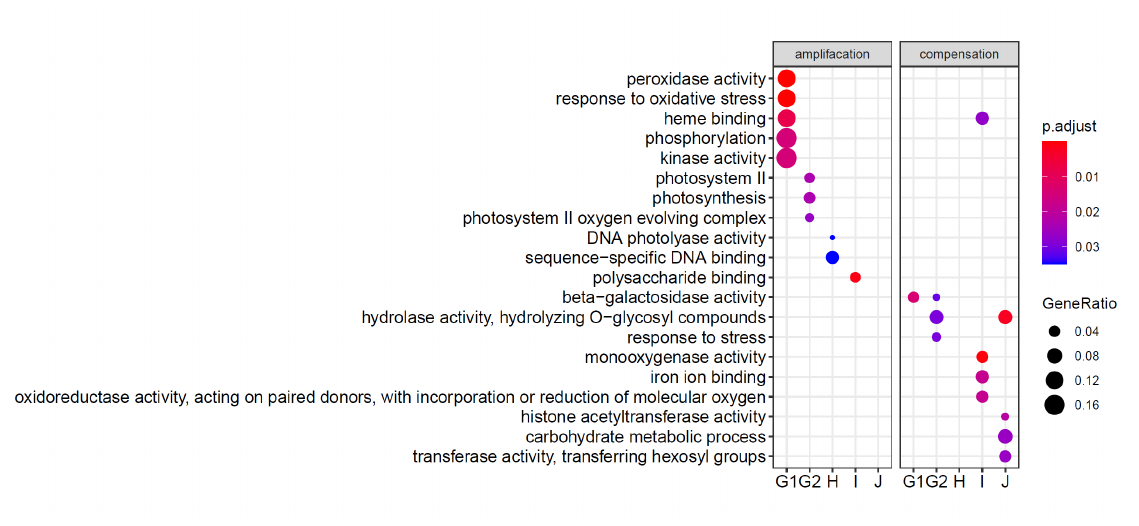

**Figure S3. Enriched GO terms of genes with amplification and compensation effects in HE individuals.**

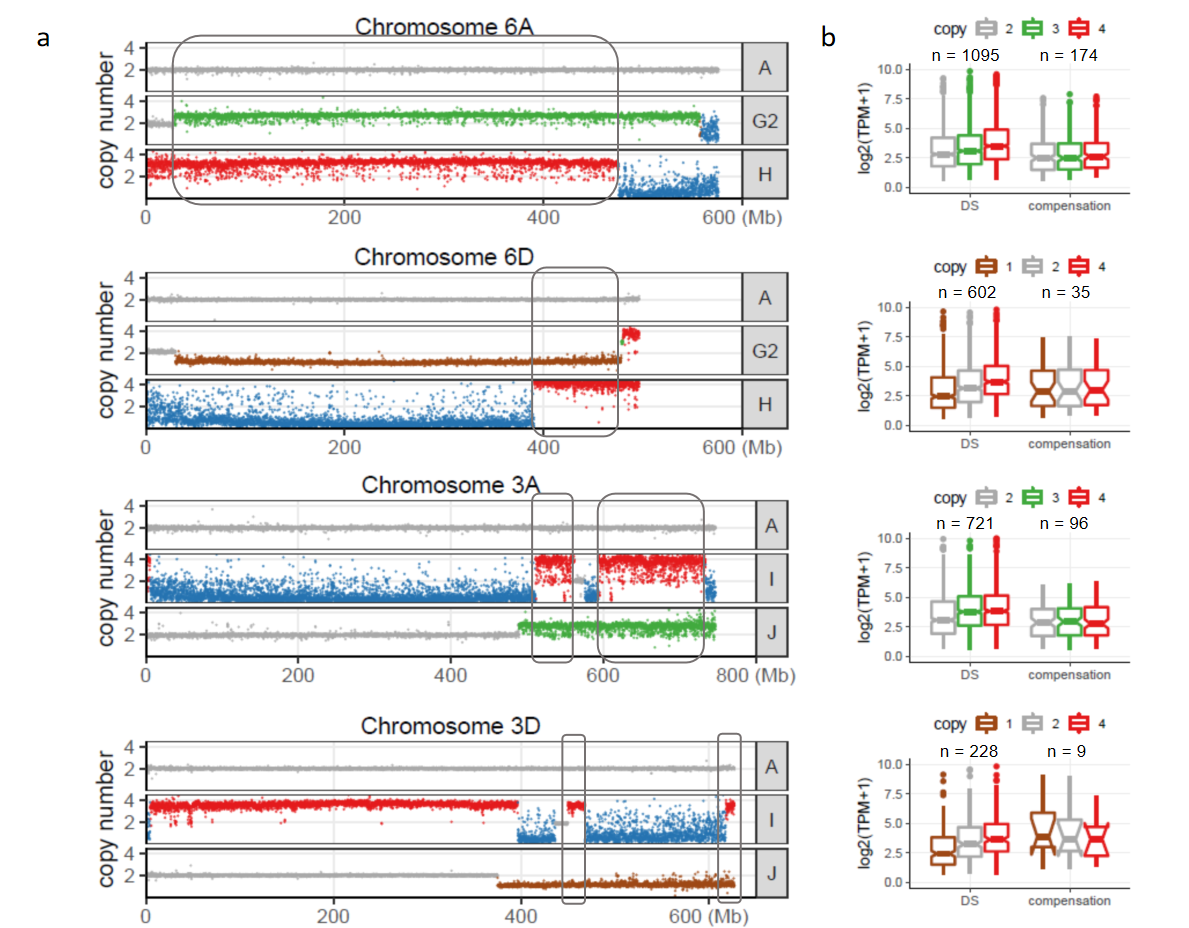

**Figure S4. Expression pattern of dosage-compensation gene in AA and DD subgenomes. (a)** Distribution of gene copy number along homoeolog chromosomes 3 and 6. Genes located in grey boxes represents they possessed different copy number among three individuals. **(b)** Expression pattens of dosage sensitive (DS) and dosage-compensation genes among variable copy numbers.

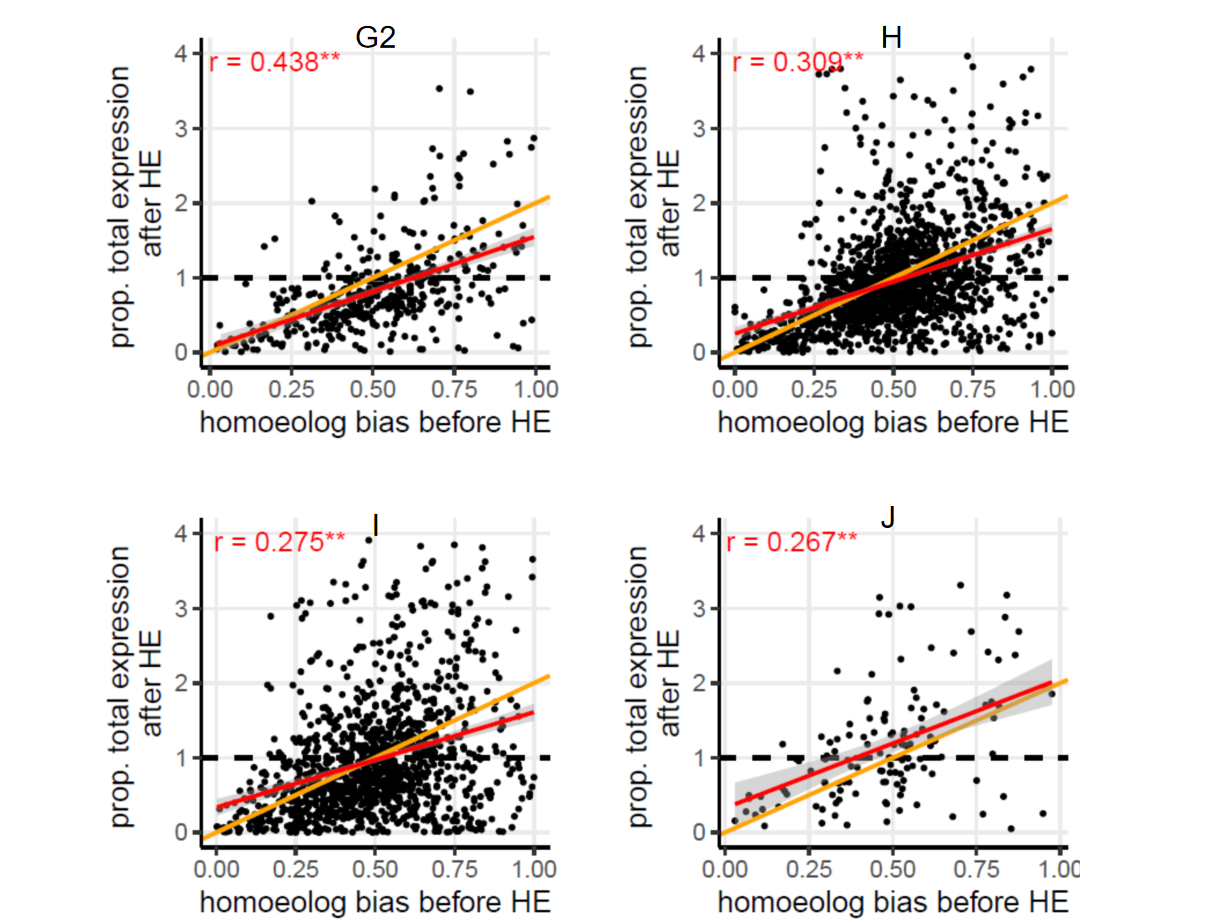

**Figure S5.** **Total expression level of homoeolog pairs (after HE) versus HEB (before HE) in HE individuals.** The x-axis represents degree of HEB and the y-axis represents the change ratio of total expression level. The black dashed line indicates the theoretical dosage-compensate pattern of total expression (y = 1). The orange line indicates the theoretical dosage-sensitive pattern of total expression (y = x). The asterisk indicates significant correlation (Pearson's product-moment correlation, p-value < 0.01). The red line fits the real data.

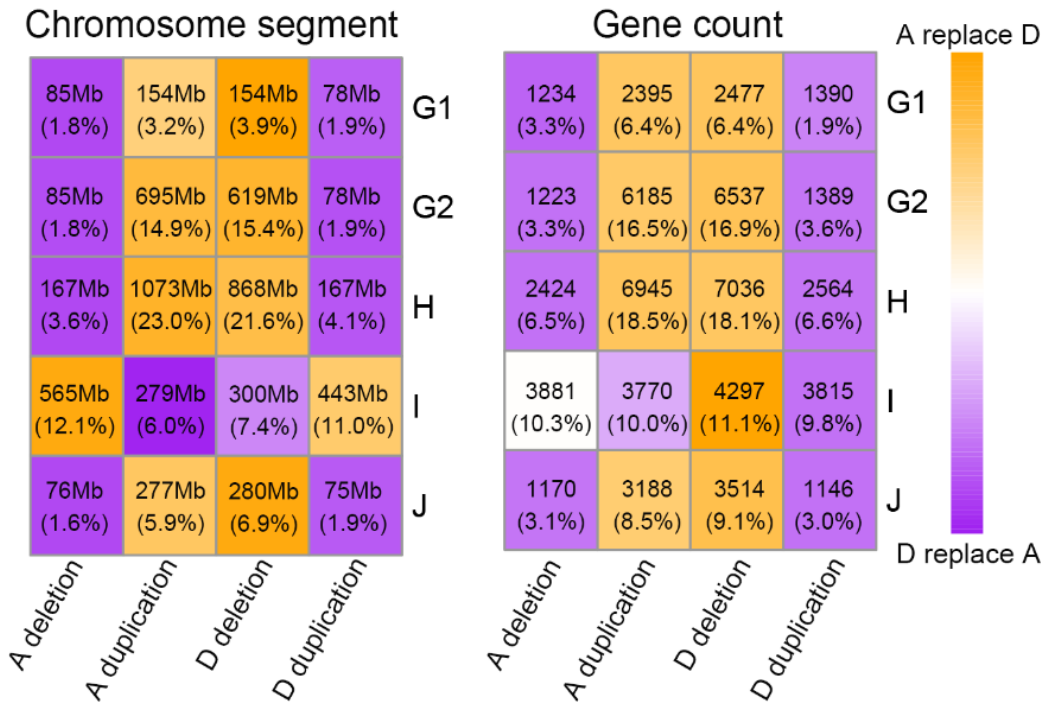

**Figure S6. Duplication and deletion information of chromosome fragment and gene count among HE individuals.**

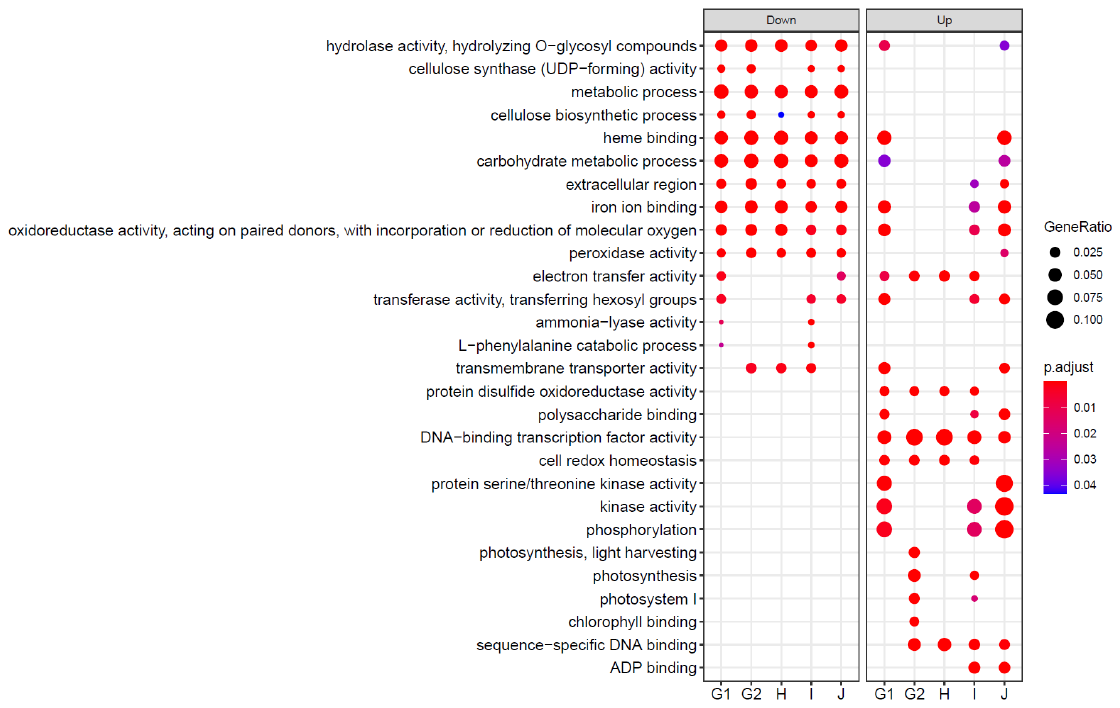

**Figure S7. GO enrichment of differentially expressed genes between control and HE individuals on non-HE chromosome segments.**

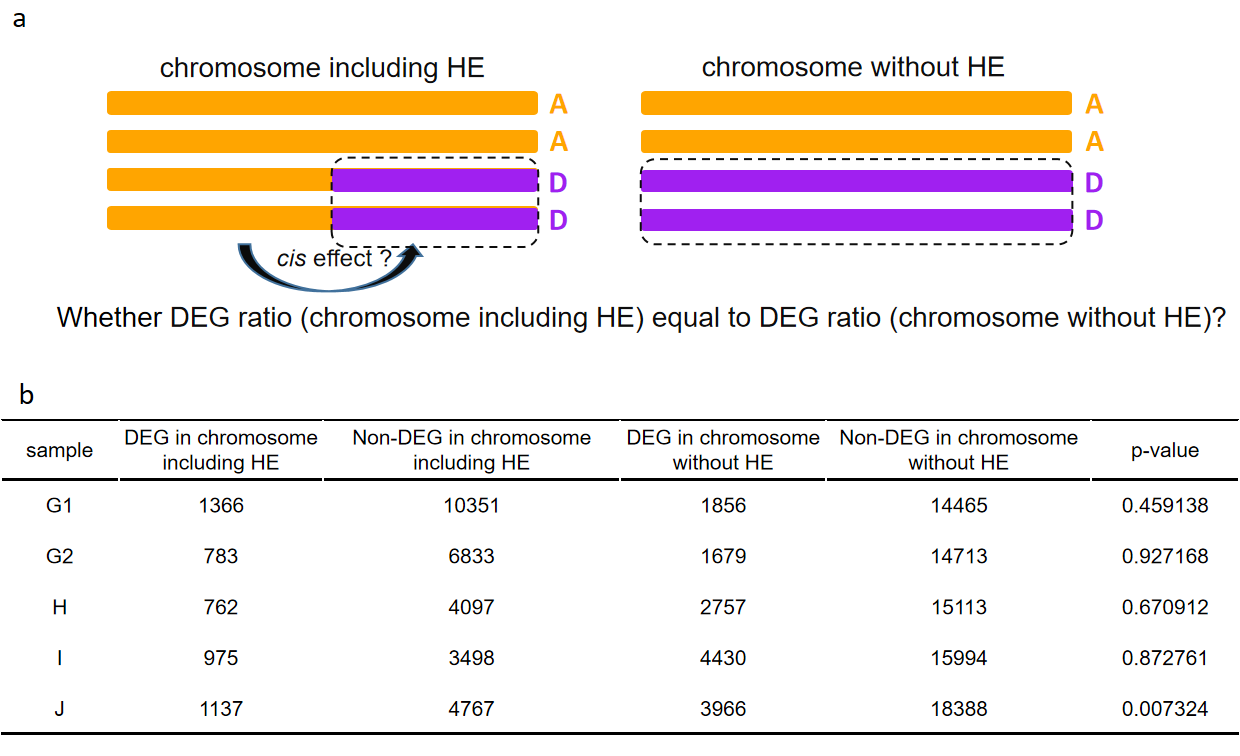

**Figure S8. Position effect (cis effect) of HE on gene expression. (a)** A schematic diagram of two classes genes with normal copies: genes located on (*i*) HE-related chromosomes and (*ii*) non-HE-related chromosomes. **(b)** Comparison of the proportion differences between two classes genes through Fisher’s exact test.

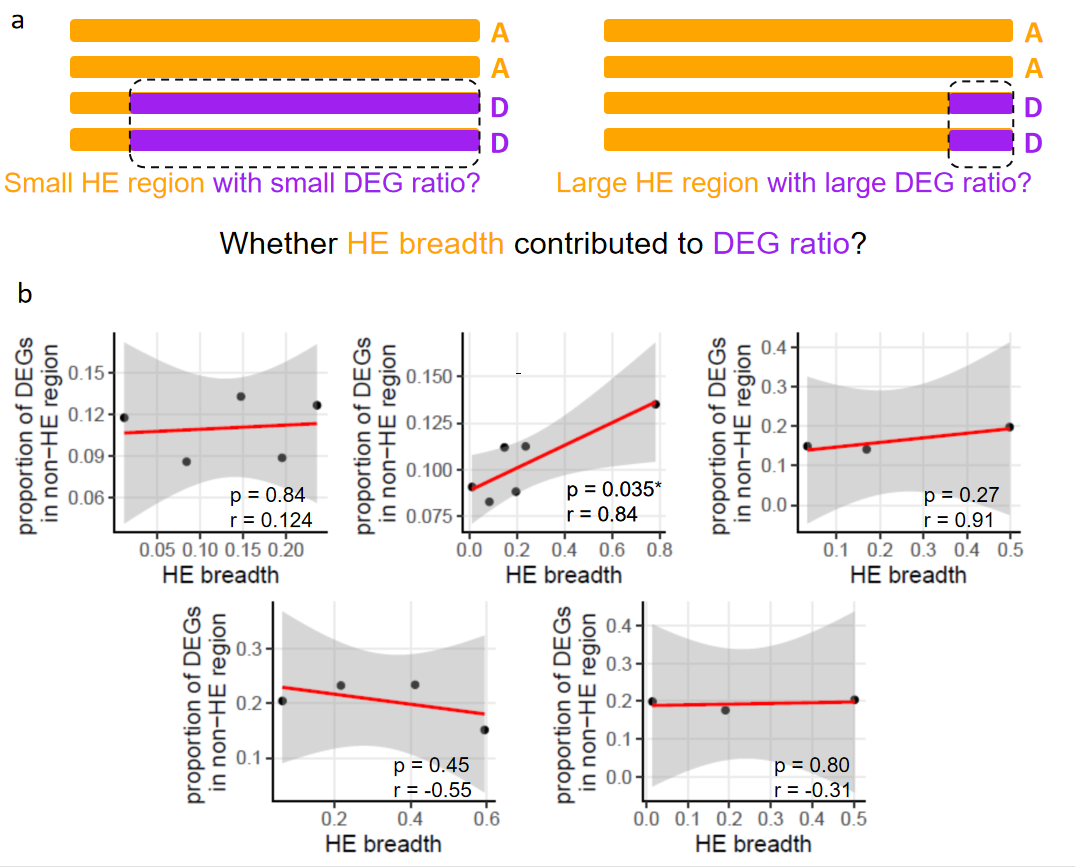

**Figure S9. Contribution of HE breadth on gene expression. (a)** A schematic diagram of two classes genes with normal copies: genes located on chromosome with (*left*) low HE breadth and (*right*) High HE breadth. **(b)** Relationship between proportion of HE-induced DEGs and HE breadth through Pearson's product-moment correlation.

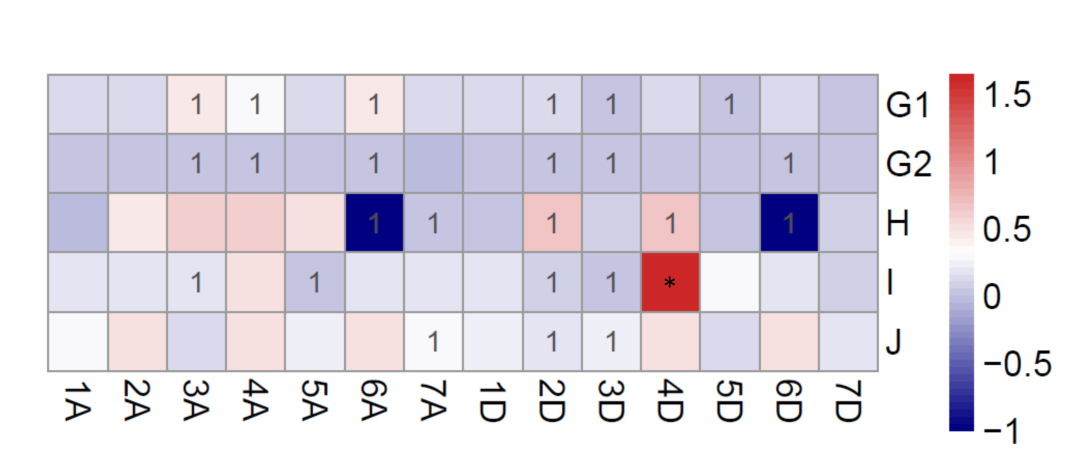

**Figure S10. Gene expression response to HE in non-HEs of different chromosomes.** Each cell represents the enrichment (red) and deficiency (blue) of DEG proportion by fisher’s exact test results. Cells marked as “1” represents HE-related chromosomes. Cells marked as “*” represents the significant enrichment od DEG proportion.

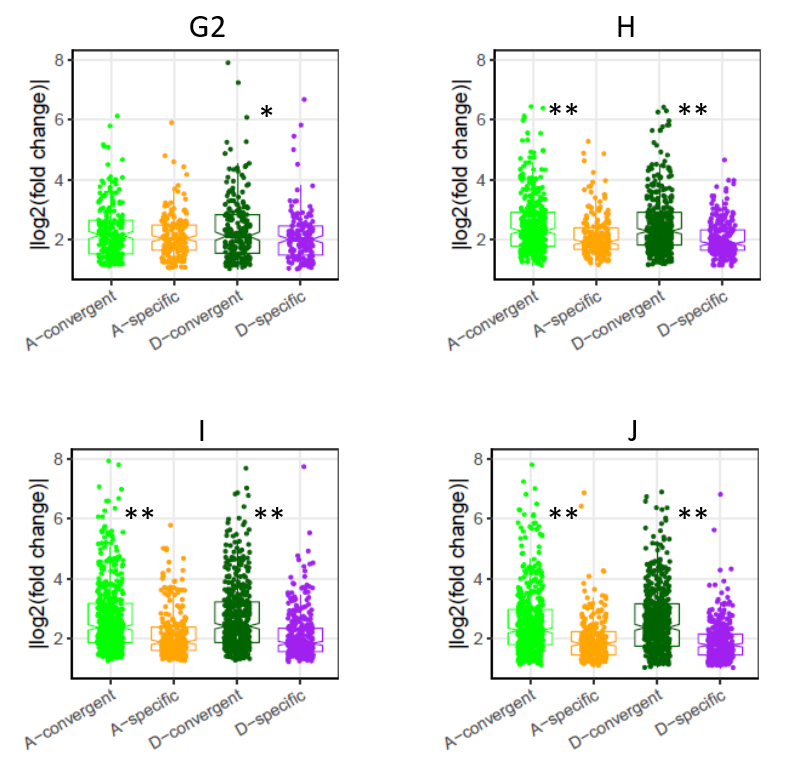

**Figure S11. Comparison of expression divergence between homoeologs in convergent and subgenome-specific classes of both AA and DD sub-genomes in HE samples.** Y-axis represents logarithm-transformed absolute value of expression ratio between control and HE individuals. The asterisks represent significant difference of expression ratio between convergent and sub-genome-specific homoeologs in AA and DD sub-genomes, respectively (Kolmogorov-Smirnov test, p-value < 0.05).

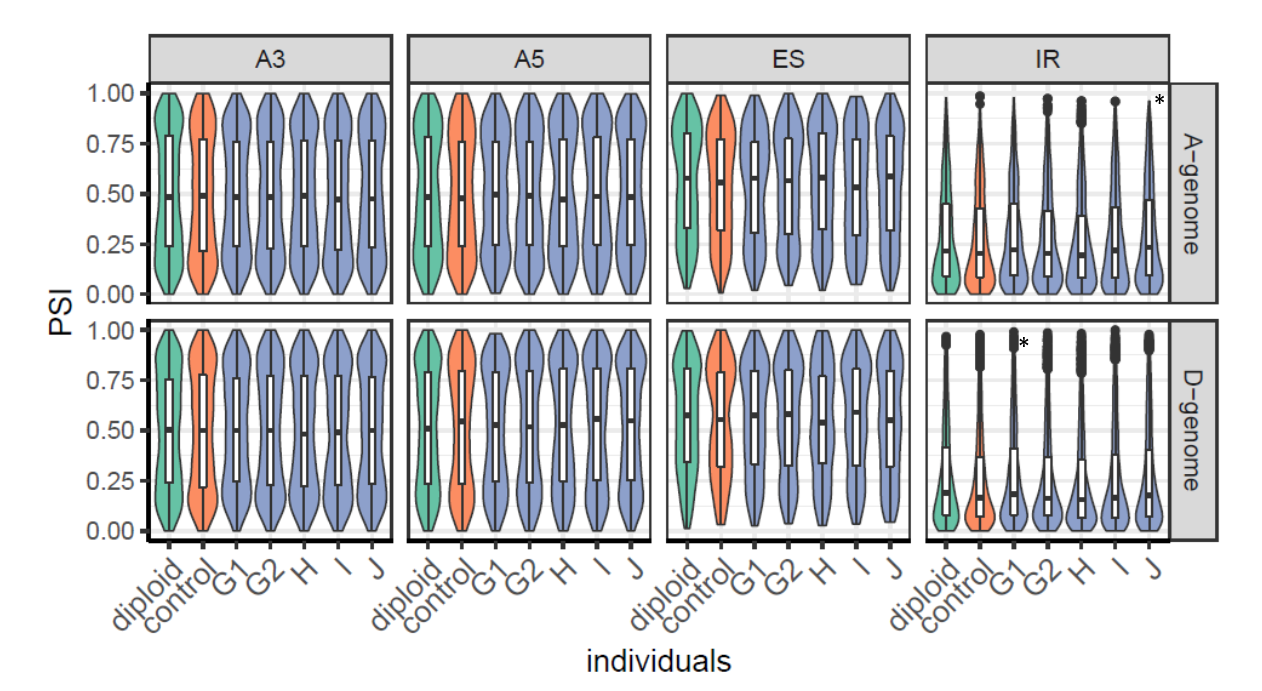

**Figure S12. ASE pattern among different types of individuals.** PSI level in diploids (green), control (orange) and HE individuals (blue) are shown. “*” represents the median PSI level of corresponding individuals is significantly higher than control tetraploid.

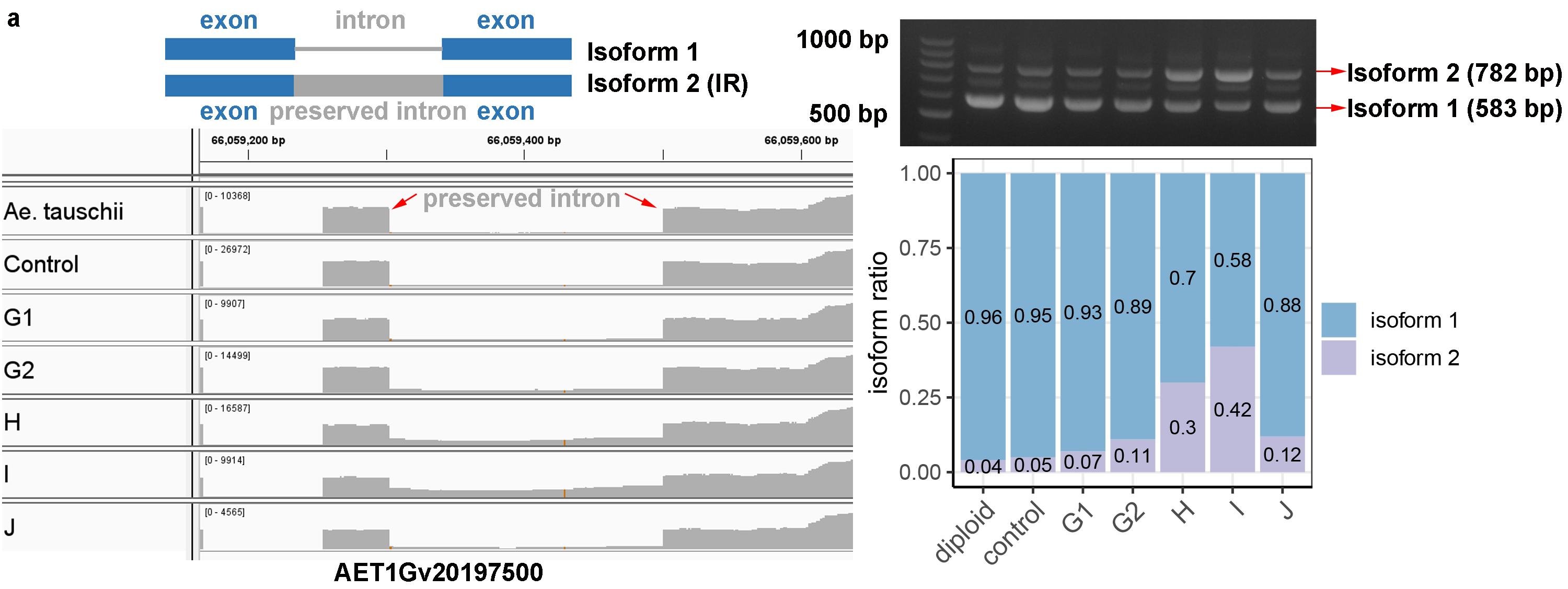

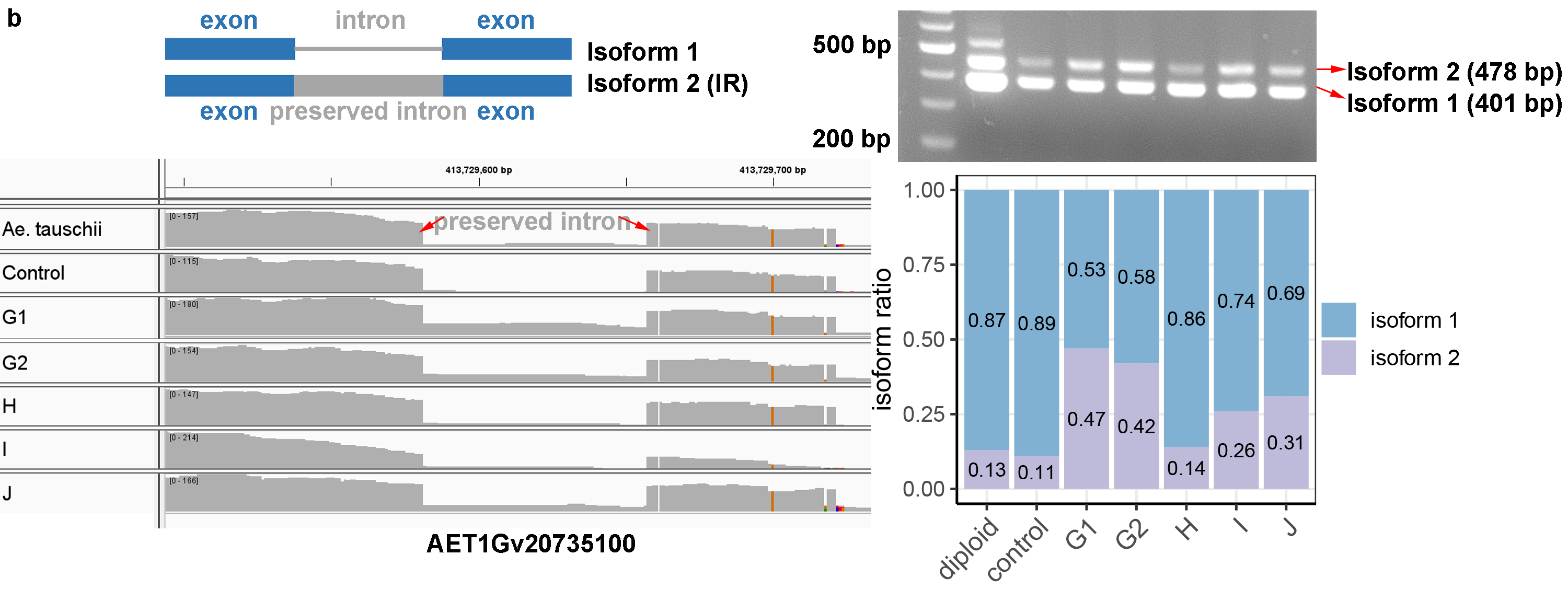

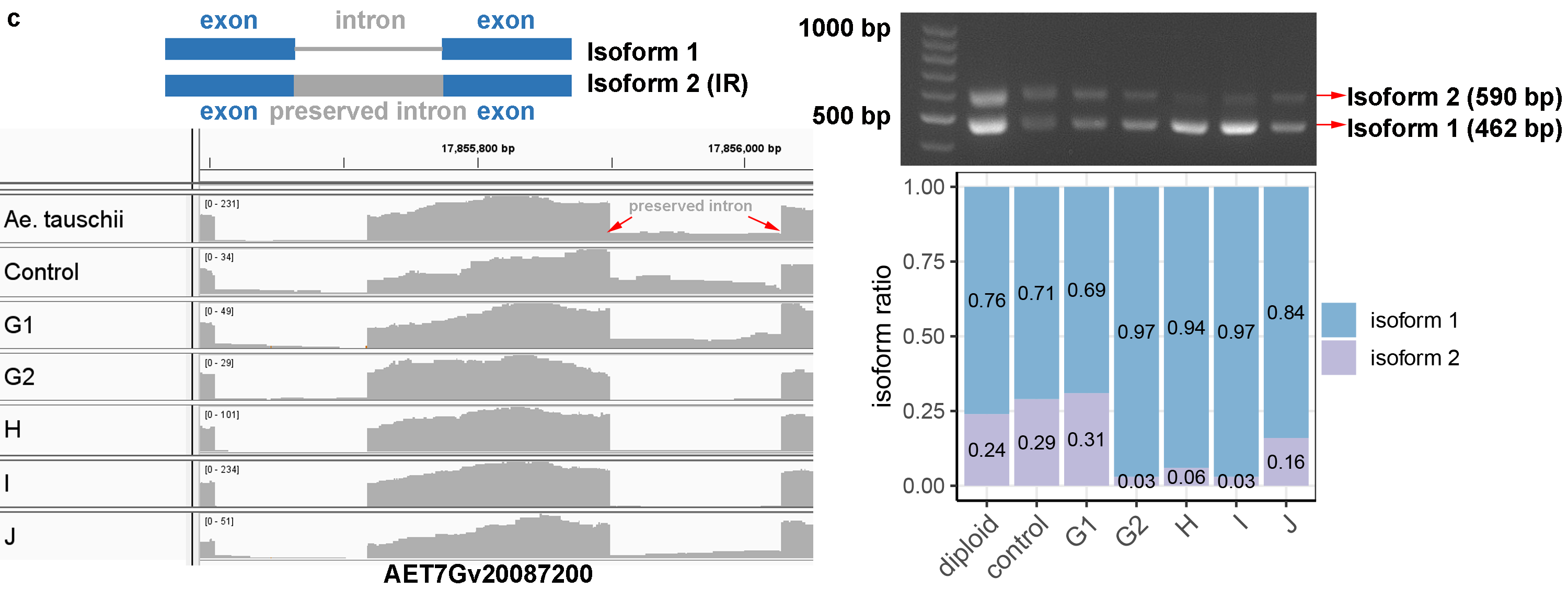

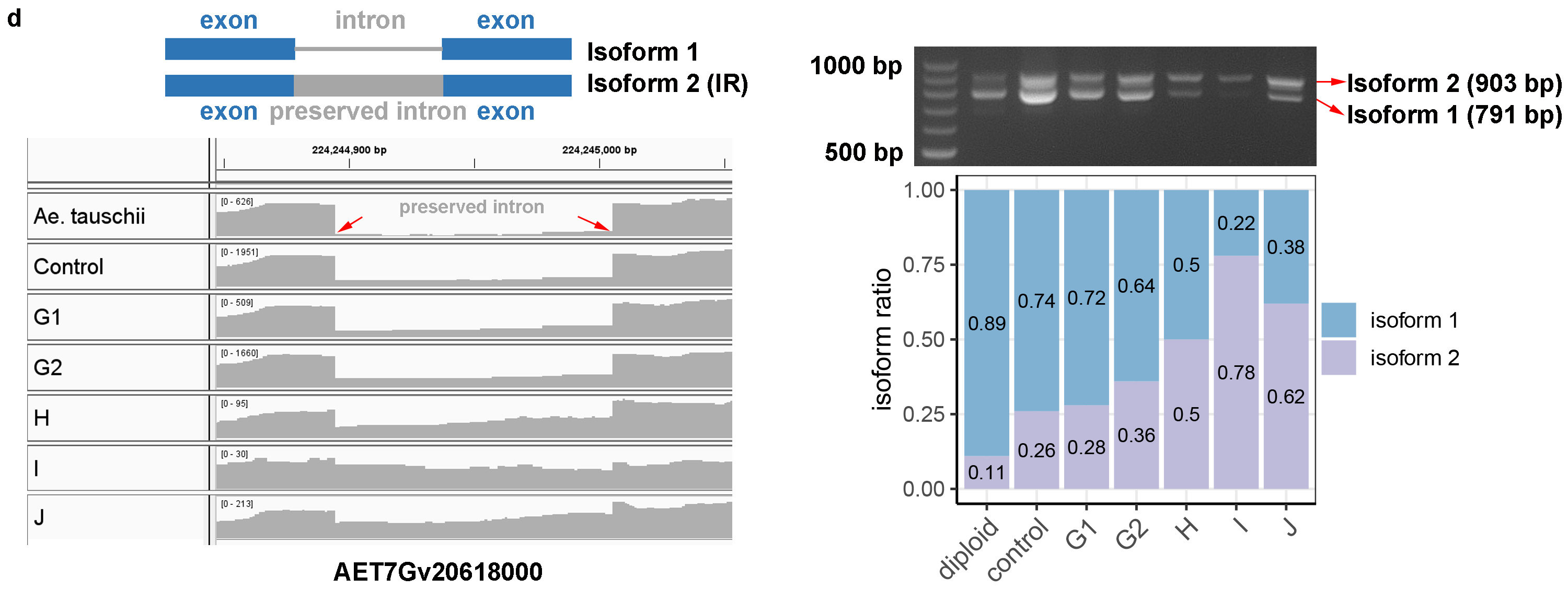

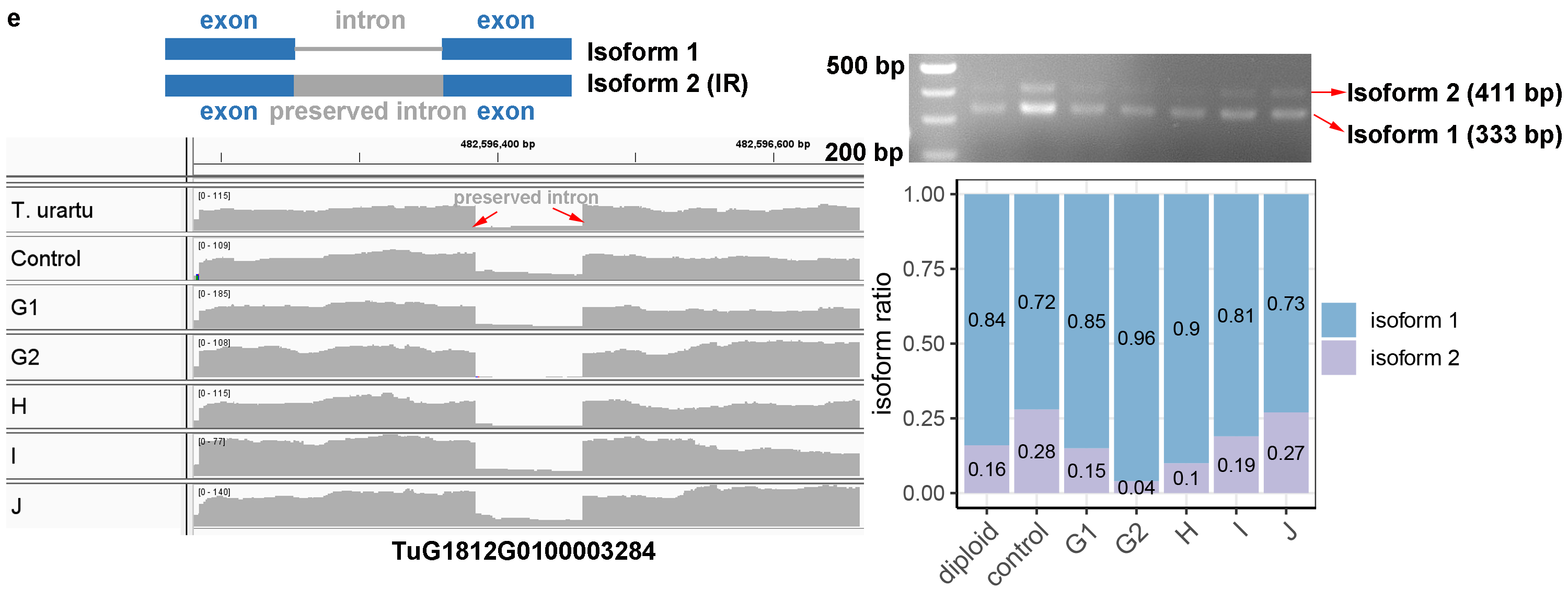

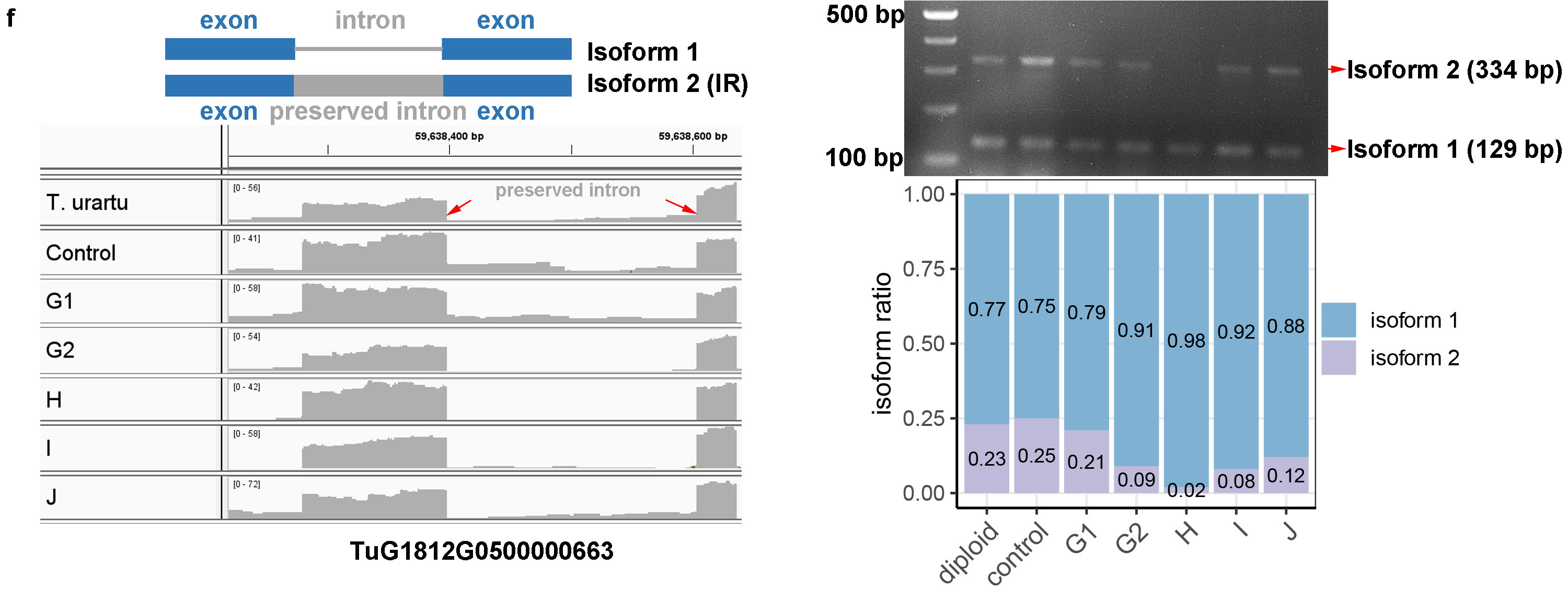

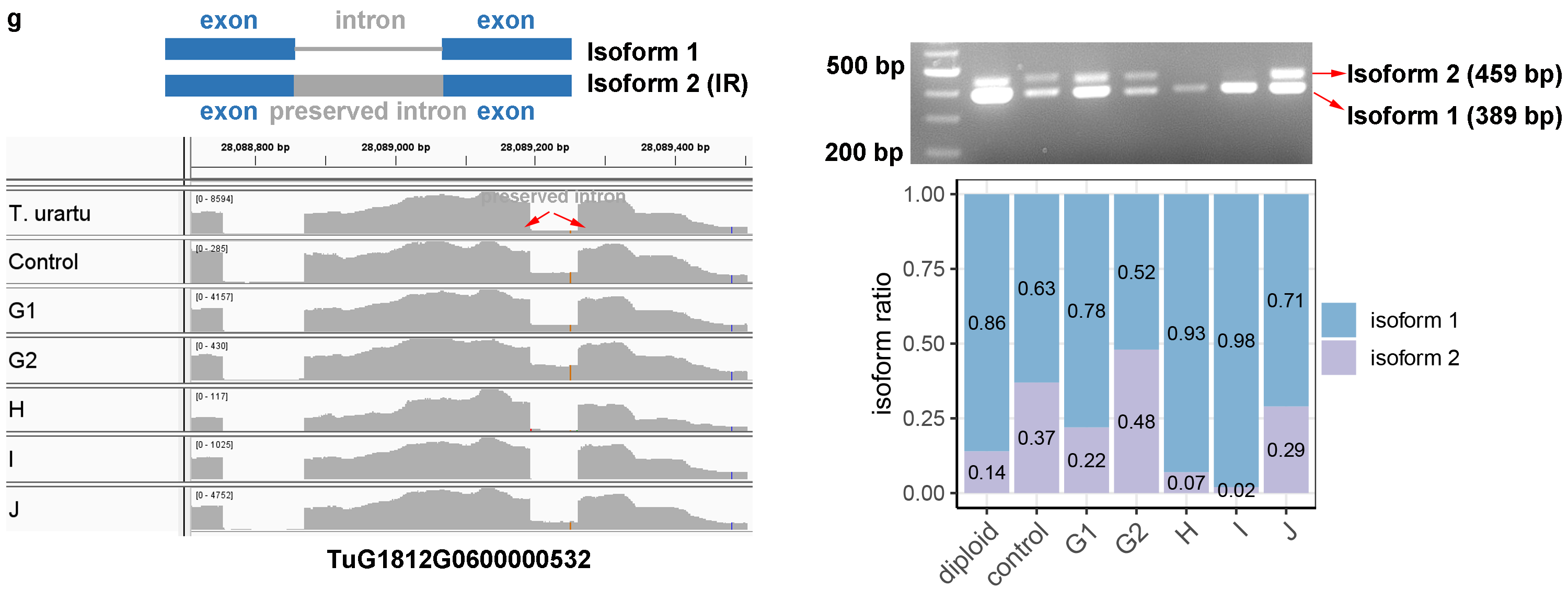

**Figure S13. Validation of ASEs between HE-containing individuals and control plant.** For each of sub-figures (**a-g**): the upper left panel shows the demo of IR-related isoforms; the upper right and lower left panel show the gal plot and IGV tracks (from RNA-seq data) of two IR-related isoforms, respectively, among diploid, control and HE-containing individuals; the lower right panel shows the relative ratios of two IR-related isoforms according to RNA-seq data.

**Table S1. Statistics of DEGs on non-HE regions between two subgenomes**

| DEG_A | None_DEG_A | DEG_D | None_DEG_D | ratio_A | ratio_D | p-value | individual |
| --- | --- | --- | --- | --- | --- | --- | --- |
| 1375 | 12018 | 1847 | 12798 | 0.102666 | 0.126118 | 7.62E-10 | G1 |
| 1100 | 10412 | 1362 | 11134 | 0.095552 | 0.108995 | 6.07E-04 | G2 |
| 1510 | 9316 | 2009 | 9894 | 0.139479 | 0.168781 | 1.07E-09 | H |
| 2417 | 9468 | 2988 | 10024 | 0.203366 | 0.229634 | 5.23E-07 | I |
| 2230 | 11247 | 2873 | 11908 | 0.165467 | 0.194371 | 2.91E-10 | J |

**Table S2. Statistics of up-regulated and down-regulated DEGs on non-HE regions**

| Down | None | Up | Up_ratio | Down_ratio | p-value | individual |
| --- | --- | --- | --- | --- | --- | --- |
| 2059 | 24816 | 1163 | 0.041479 | 0.073436 | 1.00E-56 | G1 |
| 1384 | 21546 | 1078 | 0.044902 | 0.057647 | 7.53E-10 | G2 |
| 2055 | 19210 | 1464 | 0.064411 | 0.090413 | 2.08E-23 | H |
| 3205 | 19492 | 2200 | 0.088364 | 0.12873 | 1.07E-42 | I |
| 2768 | 23155 | 2335 | 0.082631 | 0.097955 | 1.44E-09 | J |

**Table S3. Statistics of AS events in different individuals**

| type | *T.urartu* | *Ae.tauschii* | control_A | control_D | G59_A | G59_D | G63_A | G63_D | H_A | H_D | I_A | I_D | J_A | J_D |
| --- | --- | --- | --- | --- | --- | --- | --- | --- | --- | --- | --- | --- | --- | --- |
| A3 | 26.50% | 27.10% | 26.50% | 26.70% | 26.20% | 27.10% | 26% | 27% | 26.30% | 27.20% | 26.70% | 27.20% | 26.20% | 27.10% |
| A5 | 15.90% | 16.10% | 15.50% | 16.20% | 15.90% | 16.10% | 15.90% | 16.20% | 16.20% | 16% | 16.20% | 16.30% | 16.10% | 16.30% |
| ES | 7.50% | 5.50% | 7.20% | 5.50% | 7.70% | 5.70% | 7.70% | 5.70% | 7.80% | 5.90% | 7.50% | 5.80% | 7.60% | 5.90% |
| IR | 50.10% | 51.30% | 50.80% | 51.50% | 50.10% | 51% | 50.40% | 51% | 49.60% | 51% | 49.50% | 50.70% | 50% | 50.80% |
| Total num. | 4847 | 5385 | 4857 | 5242 | 5219 | 5667 | 5054 | 5465 | 4945 | 4554 | 4465 | 5209 | 5109 | 5636 |

**Tables S4. Primer sequences for validation of ASEs between HE-containing individuals and control plant.**

| **Event id** | **5’-primer** | **3’-primer** |
| --- | --- | --- |
| a | GGGGATCTGCTTCTCTTCG | CACGAGCTCATTTGCCTTAT |
| b | ACAGATCCCAAGATAGTGGT | TACTTAAGATCCCAACGACTC |
| c | TTGCCTTCAACAATAGGAGT | CAAAGATACAATTCCGTGGC |
| d | TGCACAATCTTGAGTAGGTC | CTCGATTGCTTGAGTAGGAA |
| e | ACACTAACATAGGCAGGTTT | CTCAAAATCAGTACGTCCAAC |
| f | TGGATTATTCTTTCCTGTGCT | CTTCATAGACAGATCGCCA |
| g | TCTACGTTGATCCATTCGC | CTTCTCCGGTACCATCTACG |
